# POLArIS, a versatile probe for molecular orientation, revealed actin filaments associated with microtubule asters in early embryos

**DOI:** 10.1101/2020.06.20.116178

**Authors:** Ayana Sugizaki, Keisuke Sato, Kazuyoshi Chiba, Kenta Saito, Masahiko Kawagishi, Yuri Tomabechi, Shalin B. Mehta, Hirokazu Ishii, Naoki Sakai, Mikako Shirouzu, Tomomi Tani, Sumio Terada

## Abstract

Biomolecular assemblies govern the physiology of cells. Their function often depends on the changes in molecular arrangements of constituents, both in the positions and orientations. While recent advancements of fluorescence microscopy including super-resolution microscopy have enabled us to determine the positions of fluorophores with unprecedented accuracy, monitoring orientation of fluorescently labeled molecules within living cells in real-time is challenging. Fluorescence polarization microscopy (FPM) reports the orientation of emission dipoles and is therefore a promising solution. For imaging with FPM, target proteins need labeling with fluorescent probes in a sterically constrained manner, but due to difficulties in the rational three-dimensional design of protein connection, universal method for constrained tagging with fluorophore was not available. Here we report POLArIS, a genetically encoded and versatile probe for molecular orientation imaging. Instead of using a direct tagging approach, we used a recombinant binder connected to a fluorescent protein in a sterically constrained manner and can target arbitrary biomolecules by combining with phage-display screening. As an initial test case of POLArIS, we developed POLArIS^act^, which specifically binds to F-actin in living cells. We confirmed that the orientation of F-actin can be monitored by observing cells expressing POLArIS^act^ with FPM. In living starfish early embryos expressing POLArIS^act^, we found actin filaments radially extending from centrosomes in association with microtubule asters during mitosis. By taking advantage of the genetically encoded nature, POLArIS can be used in a variety of living specimens including whole bodies of developing embryos and animals, and also expressed in a cell-type/tissue specific manner.

## Introduction

Many biomolecules function as molecular assemblies such as protein complexes, nucleic acids, and lipid bilayers. Their function often depends on the changes in molecular arrangements of their constituents, e.g. conformational changes during movement of molecular motors along cytoskeletons (1–3), rotation of F_1_-ATPases (4), bending/extending of integrins (5, 6). Molecular orientation is the key information to study the relationship between such mutual changes of molecular arrangements and their function. Fluorescence microscopy has been widely used to study the structures and the dynamics of biomolecules in living cells, but monitoring the orientation of fluorescently labeled molecules in living cells is challenging even with the latest fluorescent microscopy including super-resolution approaches. Fluorophores, such as fluorescent proteins and organic compound dyes, emit fluorescence by radiating dipoles. The light emitted from a single dipole is fully polarized along the dipole axis. Therefore, when fluorescent labels are rigidly bound to molecules, the orientation of molecules can be monitored by analyzing the polarization state of the fluorescence. Fluorescence polarization microscopy (FPM) has been used for this approach (1–9) and is especially useful for detecting the presence of orderly assembled biomolecules and changes in their arrangements (10–12). In addition, FPM can report molecular orientations at a single-molecule level even in living cells when target molecules are labeled at proper density (3, 7). Super-resolution FPM methods have also been recently reported (13–16).

Despite such promising advantages and advancements in methodology, FPM has not been widely used in biomedical research because of the difficulties in the rotationally constrained labeling of target molecules with fluorescent proteins. To monitor the molecular orientation, a target molecule and a fluorescent protein must be rigidly connected so that the orientation of the target and that of the dipole of the fluorophore are sterically fixed. For many cases, intensive screening of the linkage was required for each target molecules, since there was no universal method for constrained tagging due to difficulties in the rational three-dimensional design of protein connection. A novel, more versatile and easier approach is therefore imperative for expanding the application of FPM.

For this purpose, we have developed a versatile molecular orientation probe, named the <u>P</u>robe for <u>O</u>rientation and <u>L</u>ocalization of <u>Ar</u>bitrary Intracellular <u>S</u>tructures (POLArIS). POLArIS is a high-affinity recombinant binder rigidly connected to a fluorescent protein. The recombinant binder can be screened by phage-display to target arbitrary biomolecules of interests. Thus, POLArIS can tag arbitrary biomolecules with fluorescent proteins in a rotationally constrained manner. As a proof of principle, we developed a POLArIS that specifically binds to F-actin (POLArIS^act^) and demonstrated that POLArIS^act^ reports the orientations of actin assembly in living cells. Moreover, by observing starfish embryos expressing POLArIS^act^, we found a novel F-actin-based sub-cellular architecture that associates with the microtubule aster during mitosis, which had not been previously reported, demonstrating that POLArIS^act^ is useful for detecting ordered structures made of actin filaments in living cells.

## Results

### Constrained tagging of a recombinant binder, Adhiron, with circularly permutated superfolder GFP

Most of the successful constrained tagging approaches used the direct connection of the C-terminal α-helices of target molecules to the N-terminal 3_10_ helix of GFP (8–11). To develop POLArIS, we sought a recombinant binder protein that has an α-helix available for tagging with GFP. We found that Adhiron (the nomenclature was later changed to “affimer” (17)), a small protein (∼12 kDa) with a consensus sequence of plant-derived phytocystatins (18), has an α-helix at its N-terminus lying on top of four anti-parallel β sheets (Fig. 1A). Two variable peptide regions are inserted between anti-parallel β sheets (Fig. 1A) to create binding sites for an arbitrary molecule of interest. Phage-display screening is generally used for selecting Adhirons that specifically bind to target molecules (19, 20). We chose specific three Adhirons (Adhiron-6, 14, and 24; for simplicity, we call them Ad-A, B, and C, respectively) that were reported to bind to F-actin *in vitro* (21) as our initial test case. The dissociation constant (K_d_) of these Adhirons for F-actin was reported to be less than 0.5 μM, indicating that they bind to F-actin with a higher affinity than both Lifeact (K_d_ = 2.2 μM) (22) and UtrCH (K_d_ = 19 μM) (23). To connect the N-terminal α-helix of Adhiron to the 3_10_ helix of GFP, we used a circularly-permutated version of superfolder GFP (sfGFP) recently developed in our lab (Fig. 1A), which has the 3_10_ helix at its C-terminus and emits fluorescence as bright as the original sfGFP (9) (hereinafter simply called cpGFP). After the elimination of the end of N-terminus that does not form α-helix, exposed terminal helices of Adhiron and cpGFP were connected with a linker (L5). The sequence of L5 was reported to form an α-helix (24). We made variations of Adhiron-cpGFP fusion constructs with extended α-helix linkers (L1-4), truncated α-helix linkers (L6-9), and constructs without any linkers (L10 and 11). No serious steric hindrance (e.g. collision of main and/or side chains) between cpGFP and Adhiron connected by candidate linkers was presumed by UCSF Chimera (25) inspection for all fusion proteins (Fig. 1B shows an example for cpGFP-Ad-A L5).

**Figure 1.**
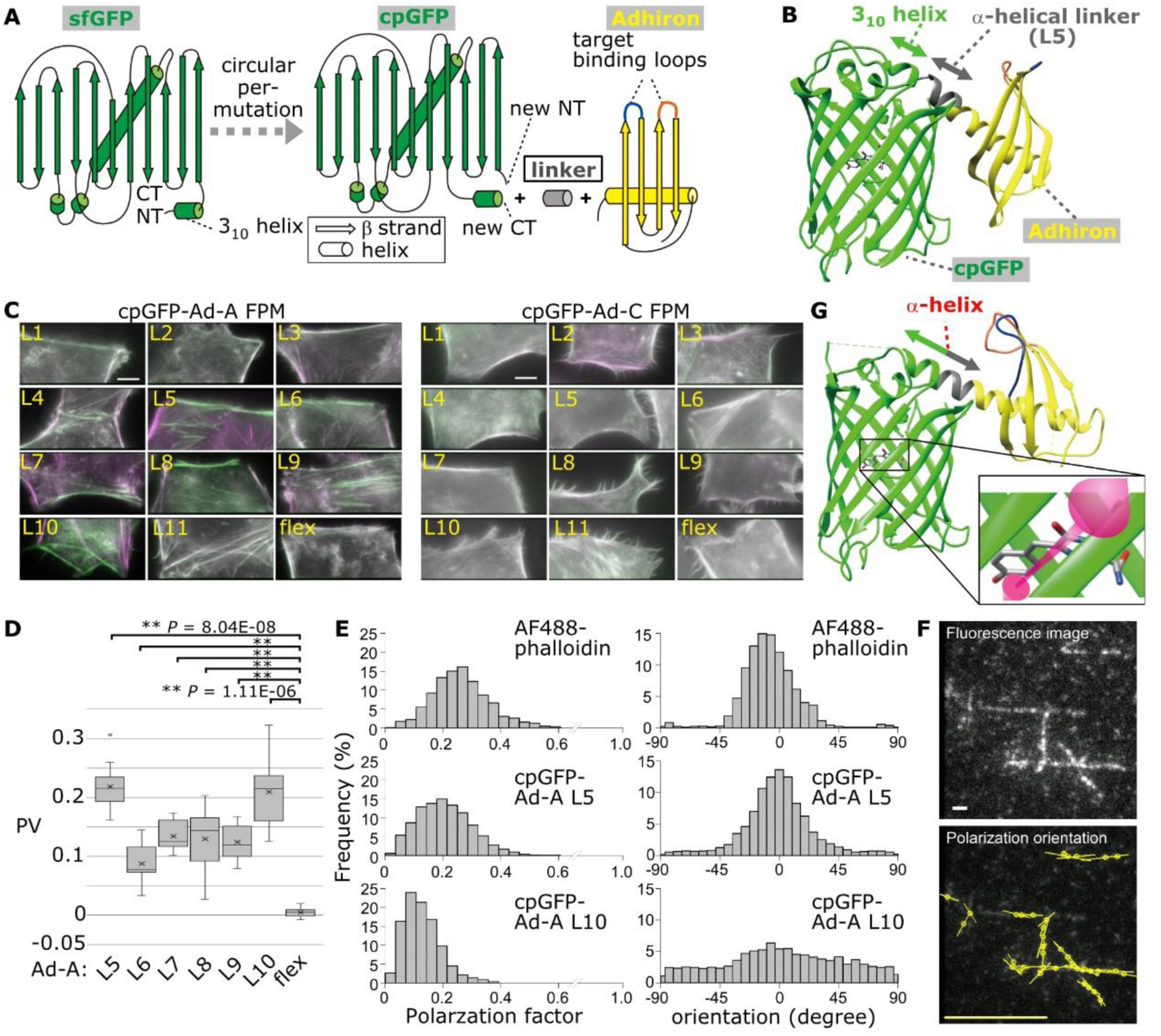
**Design of POLArIS and development of POLArIS^act^.** (A) Schematic of the circular permutation of sfGFP to create cpGFP, and fusion of cpGFP and Adhiron with a linker. (B) Modeling of cpGFP-Adhiron (L5) structure with UCSF Chimera. (C) Two-axis FPM images of HeLa cells expressing cpGFP-Ad-A/C constructs. Horizontal and vertical components are shown in green and magenta, respectively. Scale bar: 10 µm. (D) PVs of indicated cpGFP-Ad-A constructs are shown by boxplots with mean values (cross marks). Statistics: Student’s t-test with Šidák correction. **, *P* < 0.01. n=10. (E) *In vitro* evaluations of cpGFP-Ad-A L5/L10 by particle analysis with instantaneous FluoPolScope. Histograms of the polarization factor (left) and polarization orientation (right) for AF488-phalloidin (top), cpGFP-Ad-A L5 (middle) and L10 (bottom). Orientation at 0° and ±90° indicates the polarization orientation of the particle is parallel and perpendicular, respectively, to actin filaments. (F) Representative images of instantaneous FluoPolScope analysis of cpGFP-Ad-A L5. Top: fluorescence image. Bar, 1 µm. Bottom: polarization orientations. The length of bars represents the polarization factor. Yellow bar: polarization factor = 0.5. (G) The crystal structure of POLArIS^act^ (T57S). Magenta two-way arrow: presumed orientation of the dipole of cpGFP (depicted according to (79)).

### Screening and characterization of POLArIS candidates for F-actin

To test the binding affinity of Adhirons to F-actin in mammalian cells, we made Adhirons connected with cpGFP via flexible linker (flex controls). As shown in Fig. S1A, exogenously expressed flex controls for cpGFP-Ad-A and C were co-localized with F-actin labeled with phalloidin-Atto565 in fixed HeLa cells. In contrast, cpGFP-Ad-B flex showed significant background fluorescence in cytoplasm and nucleus (arrows in Fig. S1A), thus we did not test Ad-B-based constructs for further analyses. Other cpGFP-Ad-A and C constructs (L1-11) showed co-localization with F-actin stained with phalloidin-Atto565 in Hela cells, similarly to the cells expressing their corresponding flex controls (Fig. S1B).

Next, we evaluated the anisotropy of the fluorescence for cpGFP-Ad-A/C constructs in living HeLa cells expressing each construct by fluorescence polarization analysis using two-axis FPM (Fig. 1C). In this FPM setup, fluorescent molecules are excited with isotropically polarized LED illumination and the emitted fluorescence is split into two orthogonal polarization orientations, namely 0° (horizontal) and 90° (vertical) (9). For the analysis, horizontal and vertical components are pseudo-colored with green and magenta, respectively. The brightness of each pseudo-color of actin filaments represents the degree of the anisotropy of fluorescence. We found that six Ad-A-based constructs, namely L5-10, showed high brightness of green and magenta color on actin filaments running horizontally and vertically, respectively, in the image plane. The orientations of fluorescence polarization for these constructs were parallel to the axes of the actin filaments.

To quantitatively analyze the anisotropy of fluorescence of the constructs, we observed the anisotropy of fluorescence at the cortex of prometaphase-arrested HeLa M cells expressing each construct. We defined the polarization value, PV, for evaluation of the degree of anisotropy of fluorescence (see Materials and Methods and Fig. S2A for details). cpGFP-Ad-A L5 and 10 showed higher PV values (the absolute value of PV > 0.2) than those of other constructs (Figs. 1D and S2B). These results were further confirmed by the analysis using the instantaneous FluoPolScope (7), which reports the absolute 2D orientation of fluorescence polarization (Fig. S2C and D).

Next, we determined the absolute polarization orientation and the anisotropy of fluorescence (polarization factor; see a previous report (7) for more details) of single cpGFP-Ad-A L5 and L10 particles bound to F-actin *in vitro*. The probes with high polarization factor and small variance in the polarization orientation histograms are preferable for studying the 3D architectural dynamics of actin filaments, such as the actin network in lamellipodia during the retrograde flow. Fluorescence polarization of cpGFP-Ad-A/C particles sparsely bound to actin filaments was analyzed using instantaneous FluoPolScope (Fig. 1E and F). Alexa Fluor 488 Phalloidin (AF488-phalloidin) that is known to show high anisotropy of fluorescence when bound to F-actin filaments (7) was used for comparison. The polarization factor for cpGFP-Ad-A L5 was higher than that of L10, similarly to that of AF488-phalloidin. The distribution of the orientation histogram obtained for L5 was as narrow as that of AF488-phalloidin, while the orientation histogram for L10 showed broader distribution (Fig. 1E). We could observe the local orientation and the dynamics of the actin network during the retrograde flow in the lamellipodia of *Xenopus* tissue culture (XTC) cells expressing cpGFP-Ad-A L5 (movie S1). Thus, we found cpGFP-Ad-A L5 as the best orientation probe for monitoring the orientation of F-actin *in vitro* and in living cells, and hereinafter call it POLArIS^act^.

### Structure determination of POALrIS^act^

The structure of POLArIS^act^ was determined at 2.5-Å resolution by X-ray crystallography (Fig. 1G). We used T57S mutant (residue 57 in cpGFP corresponds to residue 65 in the original avGFP) of POLArIS^act^ because we could not obtain the crystal of original POLArIS^act^ protein suitable for structure determination, possibly due to the peptide-bond cleavage known as “backbone fragmentation”, during chromophore formation or maturation (26) (for details, see Fig. S3). No apparent difference was reported in the structure and orientation of chromophores of avGFP and S65T-avGFP (27), thus the T57S mutant is supposed to have the same structure as that of the original POLArIS^act^.

Remarkably, the 3_10_ helix at the C-terminus of cpGFP was reorganized to an α-helix (Fig. 1B and G). The new α-helix was continuously linked to the N-terminal α-helix of Ad-A to form a long α-helix, thereby achieving the constrained tagging. Ligand-binding loops and cpGFP were distantly located, therefore linked cpGFP is unlikely to interfere with the binding of Ad-A to actin filaments. The longitudinal axis of the chromophore was positioned almost perpendicularly to the long axis of the connecting α-helix (Fig. 1G, inset).

### POLArIS^act^ preferentially binds to F-actin in vitro and in living cells

To investigate the binding property of POLArIS^act^ for F-actin, we performed the co-sedimentation assay with F-actin. We found that POLArIS^act^ binds to F-actin with about 1:1 stoichiometry and with a dissociation constant (K_d_) of 0.32 ± 0.18 µM (Figs. 2A and S4A). Pull-down experiments indicated that POLArIS^act^ has a clear preference for F-actin to G-actin, as its binding to G-actin was undetectable in the assay (Figs. 2B and S4B).

**Figure 2.**
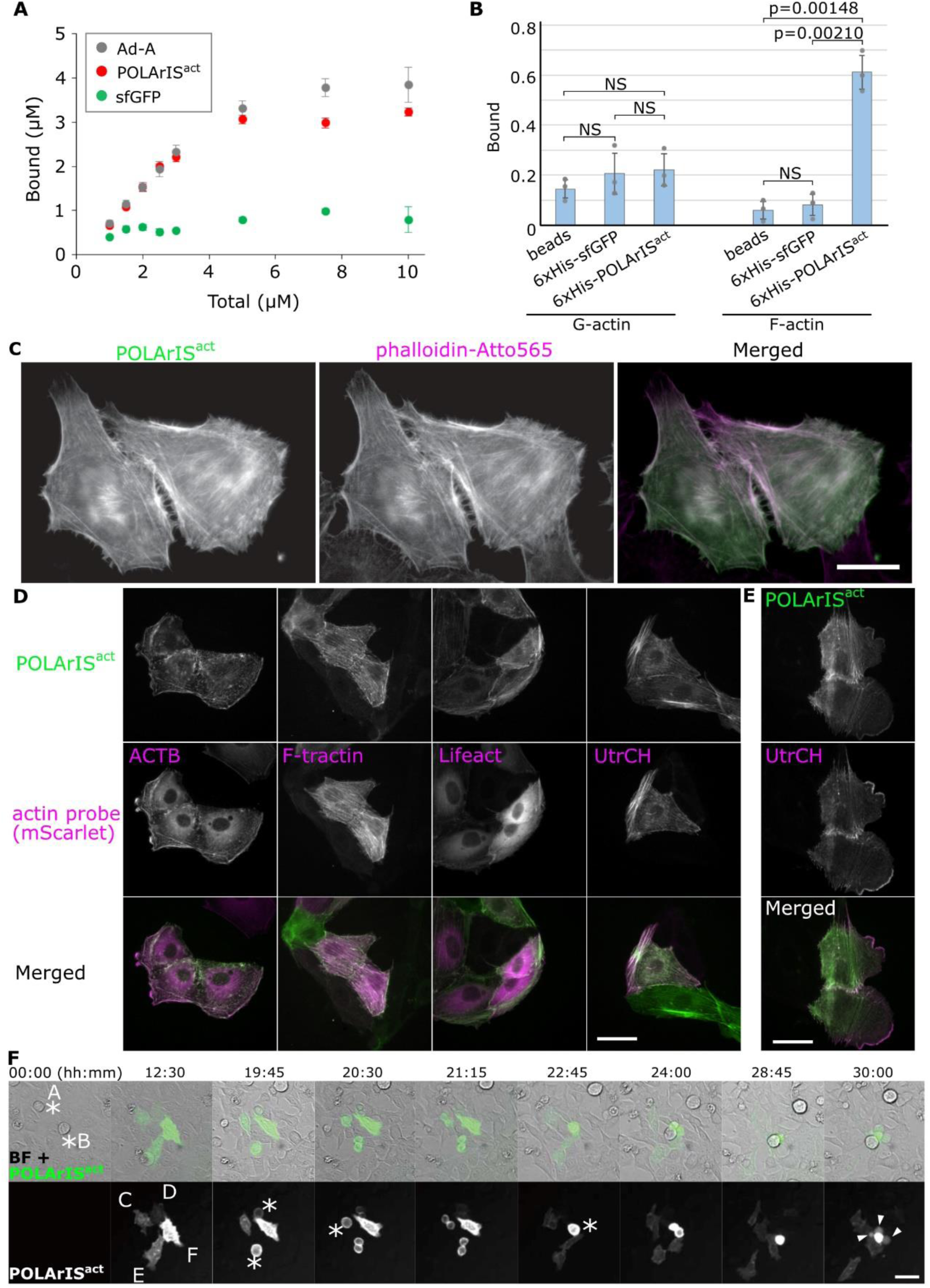
**Properties of POLArIS^act^ as an F-actin probe** (A) Plots of the mean values in co-sedimentation assays with F-actin were shown for Ad-A (gray), sfGFP (green), or POLArIS^act^ (red) with error bars showing standard deviations. Data were from three independent experiments. (B) The mean values of the bound fraction in pull-down assays were shown for G- or F-actin with the error bars showing standard deviations. *P* values were calculated by applying Šidák multiple comparison correction to Student’s t-test. Data were from three independent experiments. Gray circles indicate the values of individual experiments. (C) High-resolution images of fixed HeLa cells expressing POLArIS^act^ stained with phalloidin-Atto565. Scale bar: 20 µm. (D, E) Comparison of the localization of POLArIS^act^ with ACTB and other F-actin probes tagged with mScarlet in living LLC-PK1 cells. Compared with ACTB, F-tractin, and Lifeact, POLArIS^act^ showed a lower cytoplasmic signal (D). Localization of POLArIS^act^ was similar to that of UtrCH (D), with a relatively lower affinity to F-actin in lamellipoidia (E). Scale bars: 50 µm. (F) Live-cell time lapse-imaging of HeLa cells transfected with POLArIS^act^ plasmid. Cells A and B in mitosis at time 0 (soon after transfection) divided into cells C and D, E and F, respectively, then expressed POLArIS^act^ (time 12:30). The order of expression level was: F>E>C>D. While cells C, D, and E divided successfully, cell F failed to divide and died forming blebs (time 30:00). Asterisks indicate cells in mitosis; arrowheads indicate blebs. Scale bar: 50 µm.

The intracellular localization of POLArIS^act^ was re-investigated by co-staining with phalloidin-Atto565 in fixed cells using a high-magnification objective lens instead of a low-magnification lens we had used for the screening (Fig. S1B). The striking resemblance between their localization (Fig. 2C) proved the specific binding of POLArIS^act^ to F-actin in cells.

Next, we expressed POLArIS^act^ in living LLC-PK1 cells and compared its localization with exogenously expressed actin (ACTB), Lifeact, UtrCH, or F-tractin. The distribution of POLArIS^act^ was most similar to that of UtrCH (Fig. 2D). POLArIS^act^ showed a clearly lower cytoplasmic/nuclear background than ACTB, Lifeact, or F-tractin (Fig. 2D). As far as we have tested, POLArIS^act^ labels all F-actin structures that can be labeled by other F-actin probes, except F-actin network in lamellipodia with a moderate affinity (Fig. 2E). These results demonstrate that POLArIS^act^ can be used as an excellent F-actin marker in living cells. It is worth noting that all cell lines we tested including HeLa, HEK293, LLC-PK1, and Jurkat, grew as normally as the cells without expressing POLArIS^act^ unless POLArIS^act^ was overexpressed (Fig. 2F shows an example of HeLa).

### Fluorescence polarization analysis of F-actin in living starfish oocytes by using POLArLS^act^

We observed the fluorescence polarization of POLArlS^act^ in living starfish oocytes to test if the probe reveals assembly dynamics of actin in the physiological condition of living cells. Drastic and dynamic changes in F-actin organization have been reported during maturation and fertilization of starfish oocytes (28–33). We expressed POLArIS^act^ in starfish oocytes by microinjection of mRNA and monitored their early development processes with two-axis FPM.

First, we focused on the F-actin dynamics during the breakdown of the germinal vesicle (GV) (Fig. 3A and movie S2). Before GV breakdown, almost no fluorescence of POLArIS^act^ was observed on the GV membrane (Fig. 3A, 0 min). GV breakdown started approximately 30 min after 1-methyladenine (1-MA) treatment. POLArIS^act^-labeled F-actin abruptly appeared on the GV membrane with no apparent fluorescence polarization (5.5 min). When the GV started to shrink, we observed the fluorescence polarization signal of POLArIS^act^ as shown in the pseudo-color images of the GV membrane with green color in top/bottom regions and magenta in right/left (8 min). The transient accumulation of F-actin on the nucleoplasmic side of the GV membrane was previously observed during the initial phase of GV breakdown, called F-actin shell (28, 32). F-actin shell was reported to have two components: actin filaments running parallel to GV membrane forming a “base” of the shell, and filopodia-like spikes that protrude perpendicularly to GV membrane from the base (32). Actin filaments running in the optical plane mainly contribute to the ensemble fluorescence polarization detected by FPM. Thus, the fluorescence of POLArIS^act^ bound to actin filaments of the base is supposed to have the polarization orientation parallel to the GV membrane, while the orientation of fluorescence polarization from spikes is supposed to be vertical to the GV membrane (schematics in insets in the top row, 5.5 and 8 min). The apparent absence of fluorescence polarization (5.5 min) during the initial phase of the GV breakdown might be caused by the cancellation of fluorescence polarization originated from POLArIS^act^ bound to actin filaments in the F-actin shell (inset, 5.5 min), as the base and spikes were observed as distinct structures only with high-resolution imaging such as confocal laser scanning microscopy (CLSM) (28). The green color in top/bottom regions of GV and magenta in right/left (8 min) indicates the strong fluorescence polarization of the structures, suggesting that basal network is the dominant component of F-actin network stained with POLArIS^act^ (inset, 8 min). This is consistent with the previous reports (28, 32) showing that the actin spikes were transient structures.

**Figure 3.**
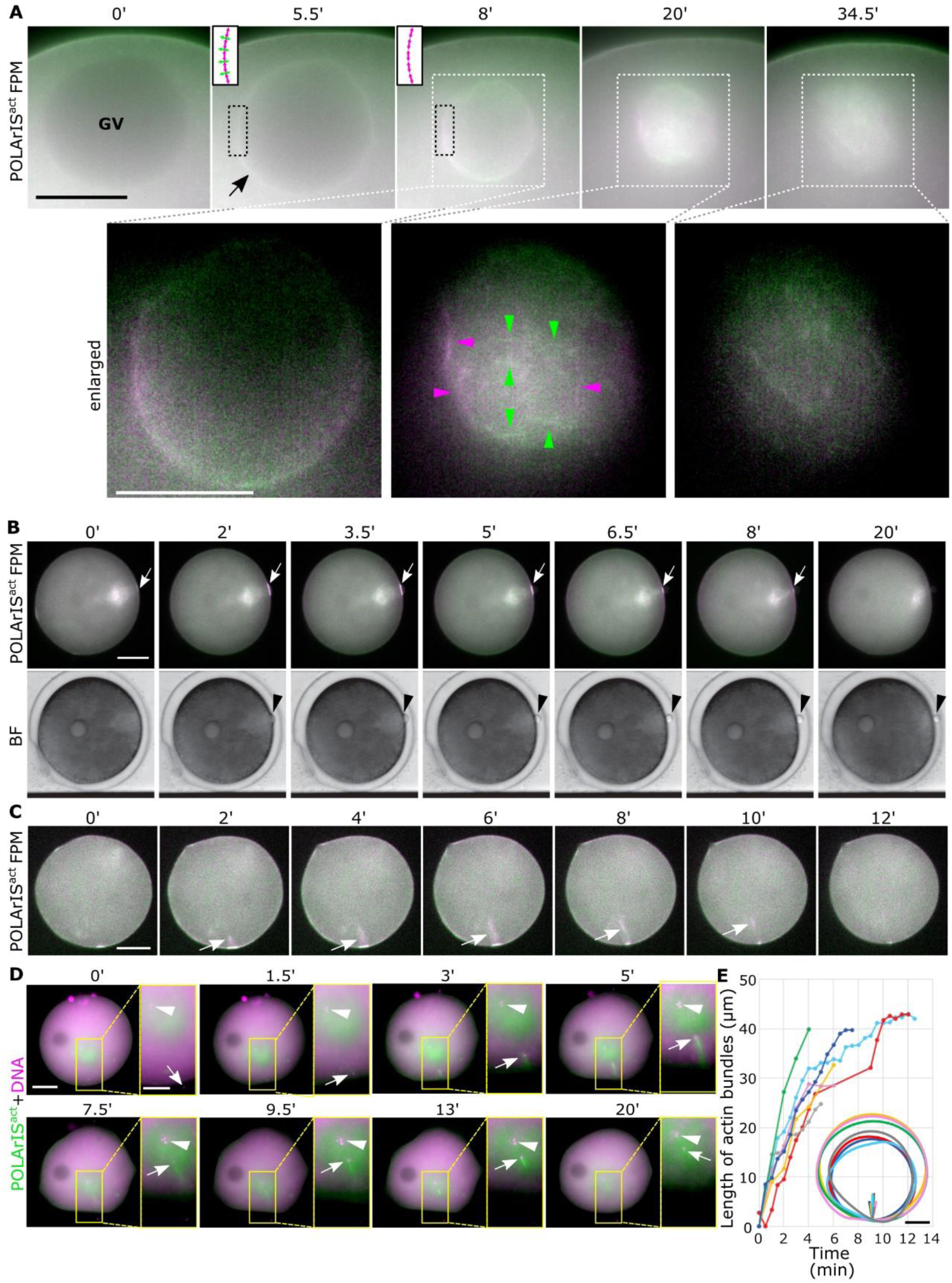
**FPM observation of GV breakdown, PB extrusion and fertilization of starfish oocytes expressing POLArIS^act^.** (A) Time-lapse images of two-axis FPM of starfish oocytes expressing POLArIS^act^ during GV breakdown. Time 0 is just before the breakdown, approximately 30 min after 1-MA addition. Regions marked by white dashed line boxes are magnified in the bottom panels (enlarged). An arrow indicates the “F-actin shell”. Arrowheads in bottom row indicate actin filaments with clear polarization in the meshwork (green: horizontal, magenta: vertical). Two-way-arrows in insets in the top row schematically indicate supposed orientations of fluorescence polarization of POLArIS^act^ bound to local F-actin structures (magenta, the base of the shell; green, spikes) in regions marked by black dashed line boxes. In bottom panels, the background signal was subtracted for the better presentation of actin meshwork. (B) Time-lapse images of starfish oocytes expressing POLArIS^act^ during the PB extrusion with two-axis FPM (top panels) and bright-field view (bottom panels). Arrowheads indicate the polar body and arrows indicate contractile ring-like F-actin, respectively. (C) Time-lapse two-axis FPM images of a starfish oocyte expressing POLArIS^act^ during fertilization. Arrows indicate actin bundles associated with fertilization. Scale bars: 25 µm (A), 50 µm (B, C). (D) Time-lapse fluorescence microscopy images of starfish oocyte expressing POLArIS^act^ (green) with Hoechst staining (magenta) during fertilization. Areas enclosed by yellow boxes in the left panels are magnified in the right panels. In some right panels, the background signal was subtracted for better visibility. After fertilization (1.5 min), sperm DNA (arrows) was positioned at the tip of the actin bundle until it stopped moving (3-13 min). The actin bundle was gradually depolymerized (9.5-13 min) and finally disappeared (20 min). Interestingly, after the extension of the actin bundle, the sperm DNA and the egg DNA (arrowheads) were closely apposed (9.5-20 min), as if the sperm DNA was conveyed to the egg DNA by the extension of the actin bundle. Scale bars: 50 µm (left), 25 µm (right). (E) Plots of the length of actin bundles over time in different colors for each oocyte (n = 8). Actin bundles typically extended perpendicularly to the cell surface and reached about 40 μm deep. The schematic shows the trajectory of the actin bundle in each egg and the outline of the same egg, with the same color as that of the plot graph. Scale bar: 50 µm.

Subsequently to the disappearance of the F-actin shell, a meshwork-like F-actin network (31, 33) was observed with fluorescence polarization of POLArIS^act^ (Fig. 3A, 20 min, arrowheads). The fluorescence signal was gradually weakened with time (34.5 min).

Following the process of GV breakdown, the polar body (PB) was extruded (Fig. 3B and movie S3). POLArIS^act^ signal began to be enriched on the cortex near the animal pole (Fig. 3B, 0-2 min, arrow). The PB appeared at the animal pole immediately after the F-actin accumulation (2 min, arrowhead). POLArIS^act^ signal rapidly decreased while half of the PB appeared on the membrane (5-6.5 min) and disappeared as the PB extrusion completed (8-20 min). In Fig. 3B, the fluorescence polarization signal of POLArIS^act^ at the PB extrusion site was clearly observed as magenta color by two-axis FPM, indicating that actin filaments are aligned parallel to the division plane. This is consistent with the proposed model of the contractile ring formation during the PB extrusion (29).

Next, we monitored the F-actin dynamics associated with fertilization (Fig. 3C-E). We found that a bright POLArIS^act^ signal appeared on the cell surface where sperm entered (Fig. 3C, 0 min). From there, an actin-based structure grew into the egg (2-8 min) similarly to the previous observations associated with fertilization (30). The strong fluorescence polarization was detected with FPM, indicating that this structure was composed of bundles of well-aligned actin filaments. The tip of the bundle, typically reaching about 40 μm from the surface membrane (Fig. 3E), had been connected to the sperm nucleus in the cytoplasm until the nucleus stopped moving in the cytoplasm (Fig. 3D and movie S4), which was reminiscent of actin-based motilities of bacterial pathogens (34). From these observations, we confirmed that our two-axis FPM setup using POLArIS^act^ can distinguish F-actin assemblies from randomly oriented F-actin accumulations in living starfish eggs and zygotes.

### Formation of radially extended actin filaments during mitosis in zygotes and blastomeres of early starfish embryos

We next observed F-actin dynamics during early development of starfish embryos. Unexpectedly, we found the strong fluorescence polarization of POLArIS^act^ observed as the green (horizontal)/magenta (vertical) cross patterns appeared in pseudo-color images during the first cleavages (Fig. 4A, second and third rows, and movie S5). The presence of the cross patterns was further examined by using the differential images of horizontal/vertical polarization (Fig. 4A, fourth row). Two small cross patterns appeared near the center of the zygote (Fig. 4A, 20 min) and rapidly expanded and reached the cell surface before the ingression occurred (24 min). The cross patterns persisted during the cleavage (39-56.5 min) and disappeared when the cleavage finished (59 min). We also observed similar structures in the blastomeres repeatedly in cleavages after the 2-cell stage (Fig. 4B and movie S6). Of note, the cross pattern also appeared using UG3, the probe of F-actin for FPM that we recently reported (Fig. S9B and C). But the pattern was not distinct probably due to UG3’s low fluorescence polarization (9).

**Figure 4.**
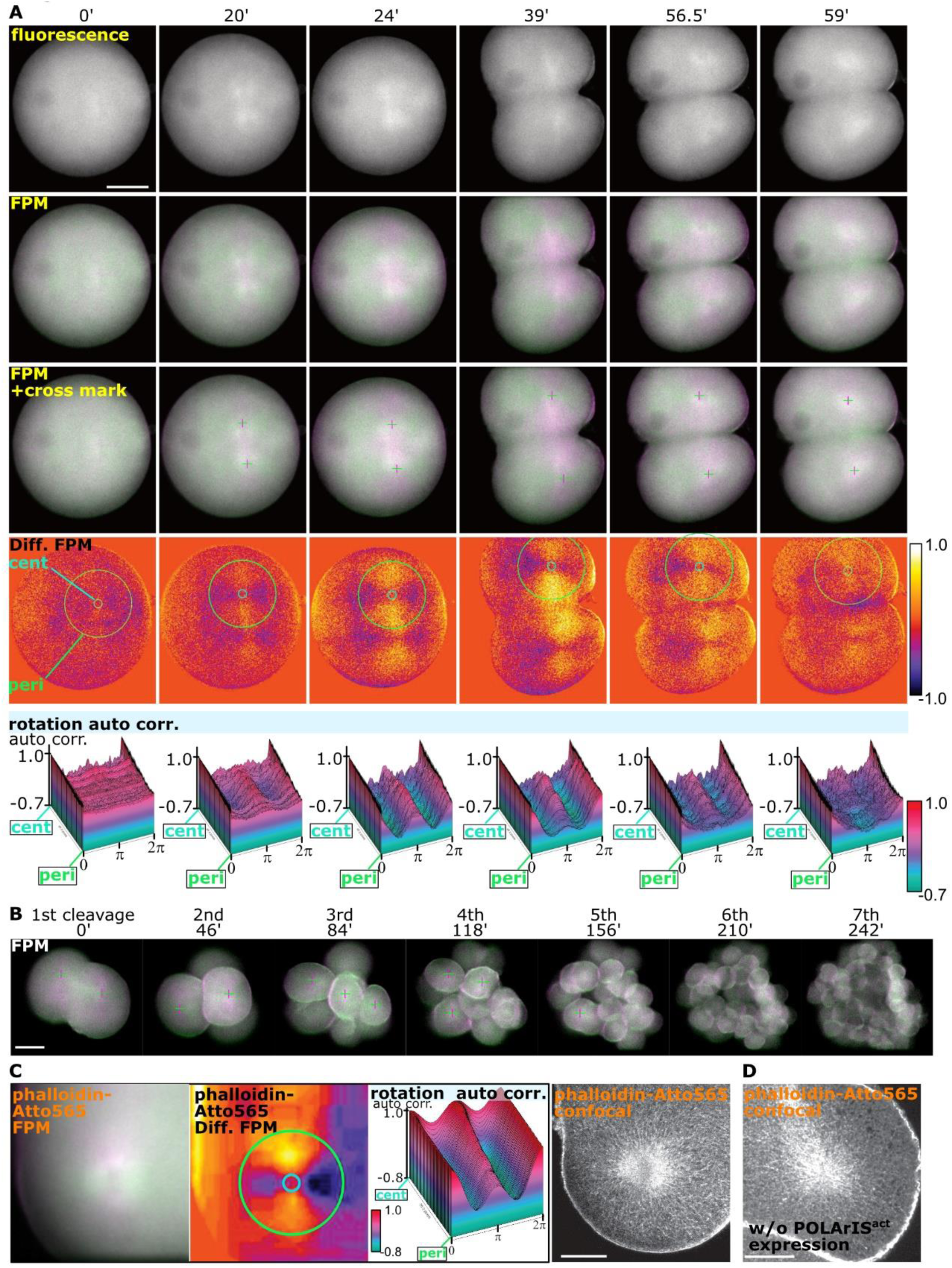
**Organization and dynamics of FLARE during starfish embryo cleavages.** (A) Time-lapse images during the first cleavage of a starfish embryo expressing POLArIS^act^ with fluorescence microscopy (the top row) and two-axis FPM (green/magenta in second and third rows and differential in the fourth row). Cross marks in the third row indicate the centers of the green/magenta cross patterns. The bottom row shows the 3D-plots of the rotational autocorrelation analyses of green/magenta cross patterns. Light blue circles (annotated as cent for center) and light green circles (annotated as peri for peripheral) in differential images indicate circles of minimum and maximum diameters used in the autocorrelation analyses. Z-axis shows the normalized autocorrelation. Scale bar: 50 µm. (B) Time-lapse two-axis green/magenta FPM observation of a starfish embryo expressing POLArIS^act^ from the first to the seventh cleavage. FLARE could not be recognized after the fifth cleavage because of the out-of-focus fluorescence from blastomeres that were overlapped with in-focus blastomeres in the Z-axis direction. Time 0 was set arbitrarily to the frame in which FLAREs were clearly visible during the first cleavage. Scale bar: 50 µm. (C) Green/magenta (left), differential (middle left, with circles of min/max diameters used in autocorrelation analyses), 3D-plot of rotational autocorrelation analysis (middle right) of two-axis FPM, and a CLSM (right) image of phalloidin-Atto565 staining of a glyoxal-fixed starfish embryo expressing POLArIS^act^ in the second cleavage. Images of FPM and CLSM were taken from the same region with almost the same z-axis planes of the same embryo. Scale bar: 25 µm. (D) CLSM image of phalloidin-Atto565 staining in a non-injected embryo during the third cleavage. Regardless of POLArIS^act^ expression, actin filaments of FLARE were observed in fixed starfish embryos. Scale bar: 25 µm.

For the quantitative analysis of the changes in the local distribution of the cross-patterns, we deduced the center of the cross pattern from the differential polarization image by image analysis, set concentric circular shells around the deduced center of the cross pattern, and plotted the autocorrelation as the circular shell is rotated around the center (see Materials and Methods for details). A cross pattern should result in high autocorrelation when rotated by π (peak) and low autocorrelation when rotated by π/2 or 3π/2 (troughs). The amplitude (the difference between the peak and the troughs) becomes higher when the cross pattern is more evident. The analyses clearly showed the dynamic appearance and disappearance of cross patterns during cleavages (Fig. 4A, bottom row, rotation auto-corr.).

The observation above implies the appearance and disappearance of actin filaments radially aligned from two centers in a wide 3D range during the cleavages. To confirm the presence of radially aligned actin filaments, we observed fixed embryos with conventional CLSM. As fixatives based on paraformaldehyde (PFA) or PFA in combination with glutaraldehyde (GA) failed to preserve these F-actin structures in dividing starfish embryos (Fig. S5), we used the recently reported glyoxal fixation (35). Although fluorescence of POLArIS^act^ was diminished by glyoxal fixation, two-axis FPM observation of phalloidin-Atto565 staining in a fixed POLArIS^act^-expressing embryo showed the characteristic cross pattern as we had observed in live starfish embryos with POLArIS^act^ (Fig. 4C, left 3 panels). Using CLSM, radially extending actin filaments were certainly visible in the same embryo (Fig. 4C, right panel). The observation of the similar radial distribution of F-actin in embryos without injection of POLArIS^act^ mRNA (Fig. 4D) excluded the possibility of artifacts caused by the expression of POLArIS^act^.

We next compared the distribution of F-actin and microtubules in a fixed starfish embryo expressing POLArIS^act^ during the first cleavage (Fig. 5A). CLSM observation revealed numerous actin filaments extended from two points. Staining microtubules with α-tubulin antibody indicated that these two points were centrosomes. Similar distributions of F-actin and microtubules were also observed during the third cleavage (Fig. 5B). Compared with microtubule-based asters, F-actin staining seemed to have a more intricate i.e. “fluffy” texture. From these observations, we named this structure as FLARE (<u>FL</u>uffy <u>A</u>nd <u>R</u>adial actin-aster associated with mitosis in <u>E</u>mbryo). To our best knowledge, no report has demonstrated the presence of this structure made of F-actin *in vivo*.

**Figure 5.**
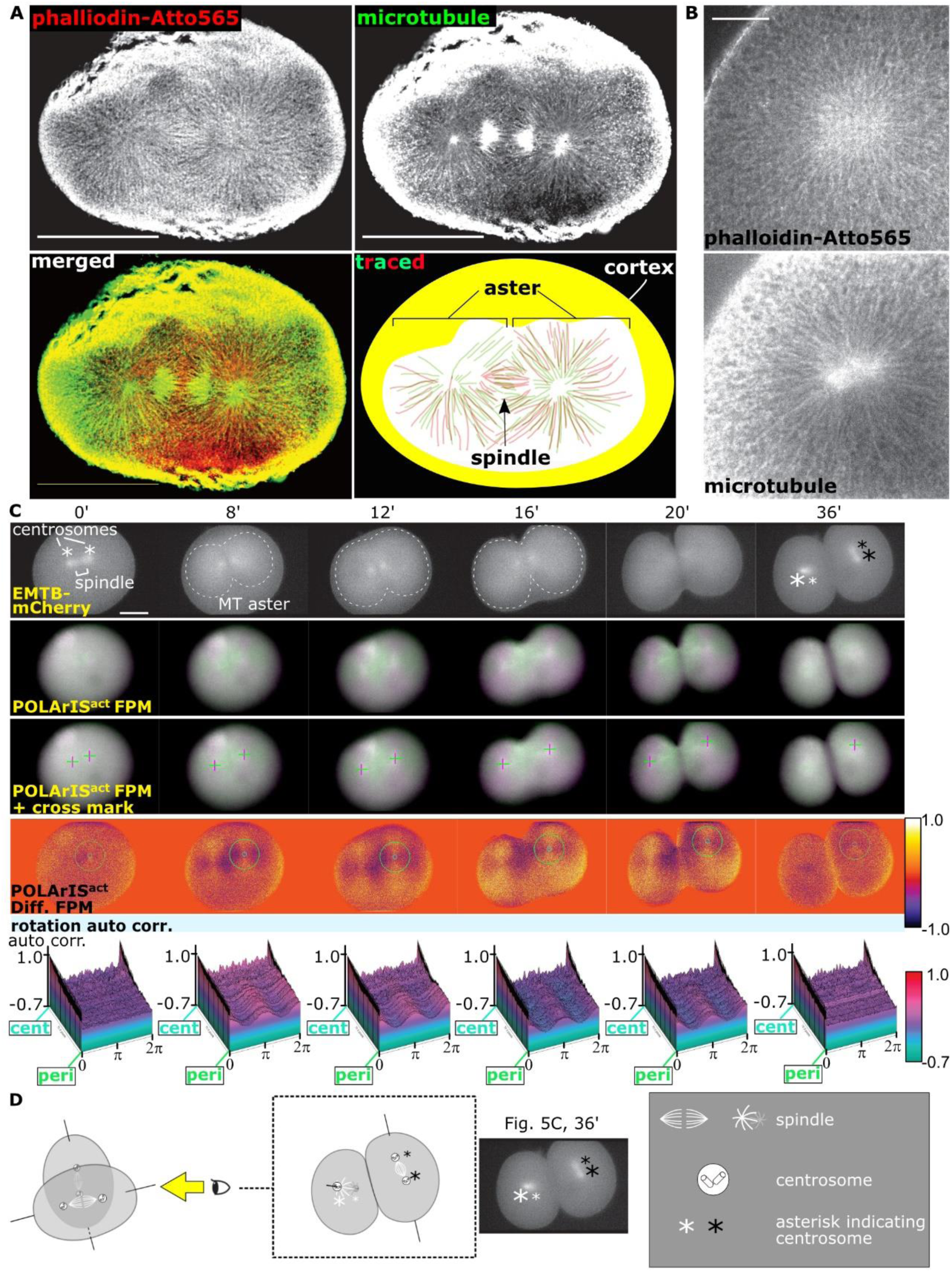
**Comparisons of the distribution and dynamics of FLARE and microtubule asters.** (A, B) CLSM images of F-actin (phalloidin-Atto565) and microtubules (anti-α-tubulin staining) in glyoxal-fixed starfish embryos expressing POLArIS^act^ in the first (A) and the third (B) cleavages. In the schematic, filaments of actin (red) and microtubule (green) were manually traced from microscopic pictures (A, bottom right). Scale bars: 100 µm (A), 20 µm (B). (C) Time-lapse green/magenta (second and third rows) and differential (fourth row, with circles of min/max diameters in autocorrelation analyses) two-axis FPM images of the first cleavage of a starfish embryo expressing POLArIS^act^ and EMTB-mCherry. The top row shows vertical polarization images of EMTB-mCherry. Cross marks (third row) indicate centers of green/magenta cross patterns. The bottom row shows 3D-plots of autocorrelation analyses of FLARE. Centrosomes are shown by asterisks (see (D) for details). Contours of microtubule asters are shown by dashed lines. In POLArIS^act^ images, the background signal was subtracted for a better presentation of cross patterns. Scale bar: 50 µm. (D) Schematic of the arrangement of spindles and centrosomes in the panel of (C), 36 min. Larger asterisks indicate centrosomes nearer the objective lens. Asterisks with the same color indicate centrosomes in the same blastomere.

### Dynamics of FLARE is tightly associated with that of microtubule aster

We next carried out live-cell observation of FLARE with microtubules visualized with mCherry-tagged Enconsin microtubule-binding domain (EMTB-mCherry) (36) (Fig. 5C and movie S7). With EMTB-mCherry, microtubule asters were observed as widespread staining of mCherry rather than radially aligned microtubules (Fig. 5C, contours of asters were indicated by white dashed lines). FLARE and microtubule aster appeared at the same time, expanded at a similar speed, and disappeared simultaneously. Before cleavage initiation, EMTB-mCherry fluorescence was concentrated around two points corresponding to centrosomes (indicated by asterisks, see Fig. 5D for details). EMTB-mCherry fluorescence was also detected between centrosomes, which corresponds to a mitotic spindle. Very weak and tiny cross patterns of FLARE were observed around centrosomes (Fig. 5C, 0 min). Concentrated fluorescence of EMTB-mCherry gradually became faint and spread as the cross patterns of FLARE expanded (8 min). The front of the expanding EMTB-mCherry fluorescence and the cross pattern of FLARE reached cell surface at 12 min, stayed during the cleavage (16-20 min). When the division finished, mCherry fluorescence was localized only at two centers, indicating that astral microtubules were depolymerized (36 min). Similarly, the cross patterns of FLARE became very weak and small at this time point. These synchronized dynamics of microtubules and F-actin were also observed in the later cleavages of the same embryo and all observed embryos co-expressing POLArIS^act^ and EMTB-mCherry that we observed (Fig. S6).

### Dynamics and maintenance of FLARE are microtubule-dependent

As described above, FLARE is very similar to microtubule asters in their dynamics and the distribution. To test possible interactions between FLARE and microtubule aster, we carried out pharmacological experiments. First, we used cytochalasin D, the actin polymerization inhibitor (Fig. S7). Treatment of a starfish embryo with cytochalasin D resulted in the loss of FLARE formation, while microtubule aster still showed the repeated dis- and re-appearance, indicating that microtubule aster dynamics is not dependent on FLARE.

Next, to examine if the microtubule aster plays a role in the formation and/or the maintenance of FLARE, we treated starfish embryos with microtubule polymerization inhibitor nocodazole (NCZ) at different time points. When NCZ was added to the embryos immediately after microtubule asters started to extend, both microtubule asters and FLARE stopped expanding (Fig. 6A, Nocodazole, 0-1 min). EMTB-mCherry fluorescence and FLARE polarization gradually disappeared (2-11 min) and the embryo failed to complete the cleavage (18-46 min). Thereafter, cleavages did not occur, and neither the microtubule aster nor FLARE reappeared in the same embryo. These observations indicate that the expansion of the microtubule aster is a prerequisite for the FLARE formation. Next, we treated an embryo with NCZ after microtubules sufficiently extended throughout the cytoplasm. In this case, microtubule aster was depolymerized outwardly from its center and the fluorescence polarization of FLARE disappeared simultaneously (Fig. 6B, 1’45”-4’45”), indicating that the maintenance of FLARE requires the integrity of astral microtubules. These results demonstrate that both the formation and the maintenance of FLARE depend on the dynamics of microtubules and the integrity of the microtubule aster.

**Figure 6.**
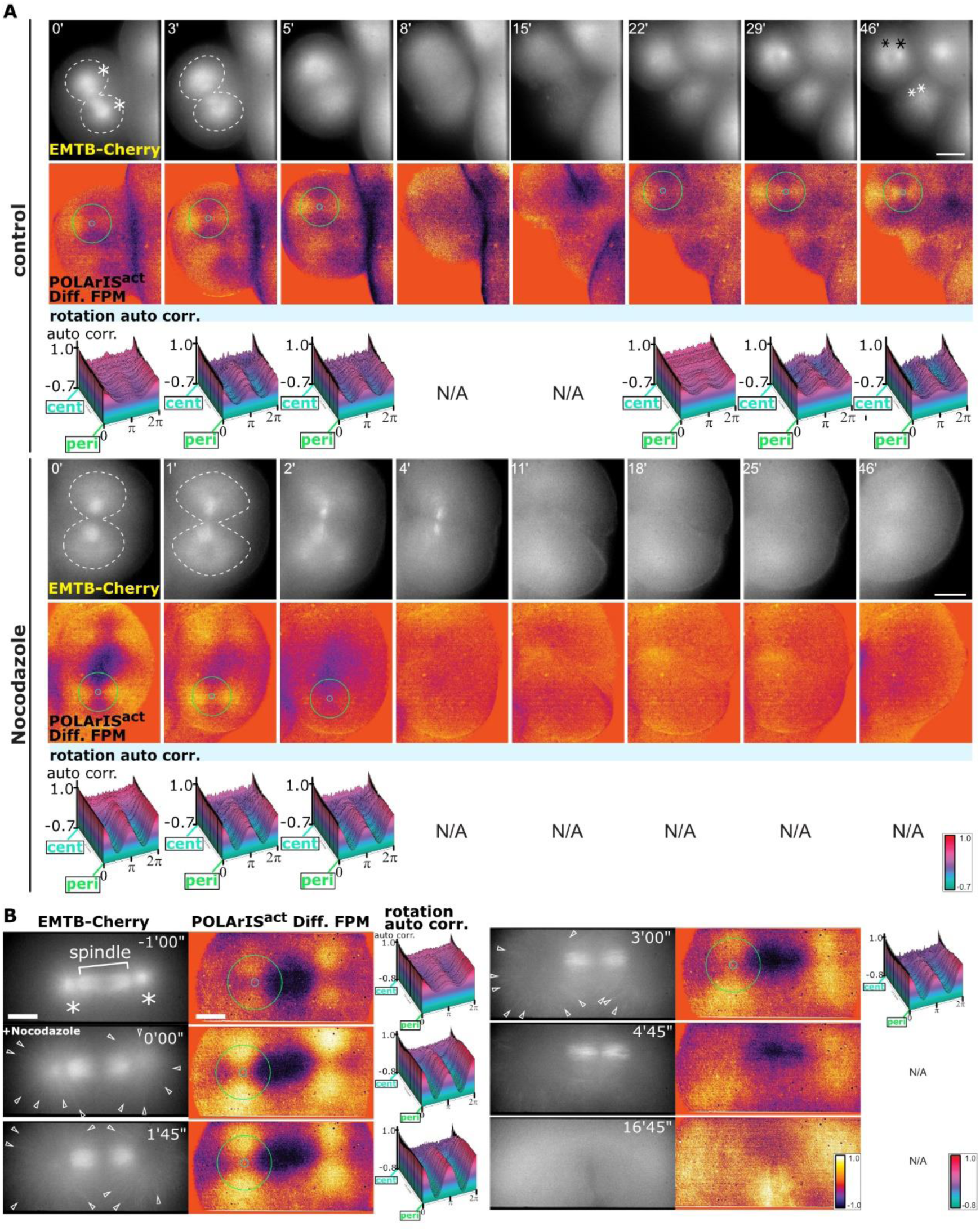
**FLARE dynamics is disrupted by inhibiting polymerization of microtubules** (A, B) Time-lapse two-axis FPM observation of DMSO (control) or nocodazole-treated embryos expressing POLArIS^act^ and EMTB-mCherry in the fourth (A) and the third cleavages (B). (A) Top rows show vertical polarization images of EMTB-mCherry (asterisks indicate centrosomes, dashed lines indicate contours of microtubule asters), middle rows show differential FPM images of POLArIS^act^ with circles of min/max diameters used in autocorrelation analyses, and bottom rows show 3D-plots of autocorrelation analyses of FLARE. N/A means that no center of the cross pattern was identified in our search algorithm. The drug was added at time 0. Scale bar: 50 µm. (B) Left columns show vertical polarization images of EMTB-mCherry (asterisks indicate centrosomes), middle columns show differential FPM images with circles of min/max diameters used in autocorrelation analyses, and right columns show 3D-plots of autocorrelation analyses of FLARE structures. N/A means that no center of the cross pattern was identified in our search algorithm. The drug was added after at least one astral microtubule reached the cell surface (time 0). A silicone immersion 40x objective lens and a 1.5x intermediate magnification lens were used to visualize microtubule filaments (arrowheads). Scale bar: 25 µm.

### FLARE formation is not Arp2/3 complex dependent

Arp2/3 protein complex and formin family proteins are known as major actin nucleation factors (37, 38). We attempted to determine which factor is responsible for the formation of FLARE by pharmacological inhibition of the activity of Arp2/3 complex and formins with CK666 (39) and SMIFH2 (40), respectively. Since SMIFH2 turned out to be impermeable to fertilization envelopes (Fig. S8A), we microinjected SMIFH2 under the elevated fertilization envelopes and perfused SMIFH2 directly to embryos (Fig. S8B). We unexpectedly found that not only actin but also microtubules lost their dynamics (10-30 min) and embryos failed to divide (40-90 min) in the presence of SMIFH2 (Fig. S8B). This implies that SMIFH2 might have nonspecific side effects in starfish embryos in addition to the specific inhibition of formin, so we could not determine if formin family proteins are involved in the formation of the FLARE in our preparations.

When embryos were treated with CK666 immediately after the first cleavage (Fig. 7), subsequent cleavages did not occur but microtubule aster showed repeated disappearance and reappearance, confirming that CK666 specifically inhibited Arp2/3- dependent actin nucleation and thereby prevented embryos from dividing without affecting microtubule dynamics. FLARE also appeared and disappeared repeatedly, indicating that Arp2/3 is not essential for the FLARE formation.

**Figure 7.**
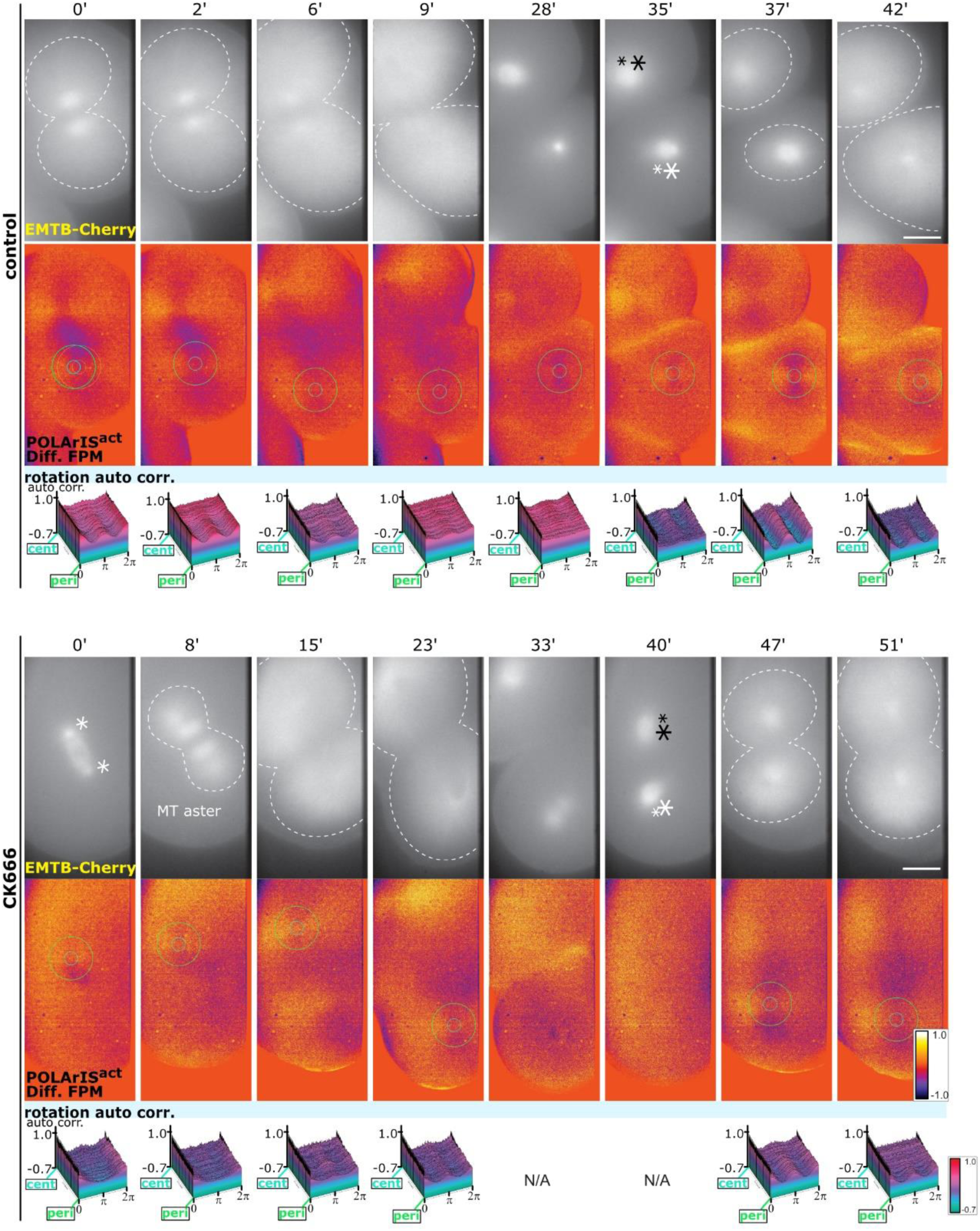
**FLARE formation and dynamics are independent of the actin nucleation by Arp2/3.** Time-lapse two-axis FPM observations of DMSO (control) or CK666 treated starfish embryos expressing POLArIS^act^ and EMTB-mCherry. Drugs were added immediately after the first cleavage completed (time 0). Top rows show vertical polarization images of EMTB-mCherry (asterisks indicate centrosomes, dashed lines indicate contours of microtubule asters), middle rows show differential FPM images of POLArIS^act^ with circles of min/max diameters used in autocorrelation analyses, and bottom rows show 3D-plots of autocorrelation analyses of FLARE. N/A means that no center of the cross pattern was identified in our search algorithm. Scale bar: 25 µm.

## Discussion

Our previous studies on the cytoskeletal dynamics (41, 42) have led us to focus on fluorescence polarization techniques, as the polarized fluorescence microscopy might detect the initial phase of polymerization even when the polymerized cytoskeletal protein molecules cannot be detected as filaments. Through the years of our trials to use FPM to analyze the dynamics of cytoskeletal proteins, we noticed that the biggest challenge was the difficulty in tagging each protein of interest with a fluorescent protein in a rotationally constrained manner. Designing more versatile and affordable methods were required for the constrained tagging of any biomolecules of interests with fluorescent labels. We have achieved this task by using a small recombinant binder protein Adhiron/affimer as a scaffold to connect target molecules with circularly permutated sfGFP. As described in this report, we established a new tool, POLArIS, taking advantage of Adhiron/affimer that can be screened through phage-display for any specific target molecules of interest. There are many Adhirons/affimers reported, targeting variety of specific biomolecules (19, 20, 43–49) from laboratories and the production service is commercially available. The two-axis fluorescence polarization imaging is a robust system for mapping fluorescence anisotropy with the aid of commonly used software for image analysis (NIS-elements Advanced Research from Nikon and ImageJ). We expect that POLArIS will help broader range of biomedical researchers to use fluorescence polarization techniques without any difficulties in the constrained tagging not only for proteins but also for nucleic acids or lipids.

Here, by using POLArIS^act^ as the first POLArIS example of POLArIS series, we show that POLArIS^act^ can report the orientation and localization of F-actin in living cells. The observations of starfish eggs and embryos expressing POLArIS^act^ revealed a previously unidentified highly ordered F-actin based architecture, FLARE. We did not find such ordered structures with conventional fluorescence microscopy, but with POLArIS^act^, radially orientated actin bundles extending from centrosomes were visible by their polarized fluorescence. This is reminiscent of Shinya Inoué’s first visualization of mitotic spindle dynamics in living cells using a polarizing microscope in the late 1940s, which could not have been achieved with conventional microscopic techniques for living cells (50, 51). While the association of actin filaments with spindle microtubules have been reported in several mammalian oocytes (52–54) and *Xenopus* embryonic epithelial cells (55, 56), to our best knowledge, our finding is the first report that demonstrates the presence of radially aligned actin filaments that extend towards the cortex in association with growing astral microtubules in living cells. Besides the use of fluorescence polarization technique, one of the most plausible reasons that FLARE has been left unidentified until today was the effect of fixatives. We found that glyoxal is one of the best fixatives for actin filaments in FLARE. Indeed, immunostaining of glyoxal-fixed neurons with anti-β-actin antibody showed a significantly brighter staining than that of PFA-fixed neurons (35). Recently, in *Xenopus* egg extract, the formation of radially aligned actin filaments around centrosomes were observed (57). It would be interesting to see if actin filaments form FLARE-like structures in eggs/embryos of *Xenopus* and other species during cleavages *in vivo*.

The actin filaments that extend from centrosomes towards cell cortex in mitosis may imply an actin-based physical connection between the cell cortex and centrosomes and/or the mitotic spindle. The mechanism of cytokinesis initiation e.g. how cleavage plane is determined and how furrow ingression is induced has been actively studied and is still one of the fundamental questions remain to be solved in cell biology. This is especially enigmatic in large cells such as eggs, since the mitotic spindle, which is believed to provide positional cues to induce contractile formation and furrowing, is far away from the cell cortex. While astral microtubules have been thought to have central roles in these pathways (58–60), roles of F-actin are currently known to be only at the cell cortex (61). Since the existence of actin filaments associated with astral microtubules has not been recognized until now, it would be fascinating to test if FLARE is involved in the signal transduction in cytokinesis initiation and/or other processes related to cytokinesis.

The mechanism of the FLARE formation is currently unknown. Centrosomes isolated from Jurkat cells formed radial F-actin array *in vitro* (62), which resembles FLARE, and actin filaments were found to be localized around centrosomes in HeLa and Jurkat cells (63, 64). Moreover, radially extended actin filaments from centrosomes were observed in HeLa cells during the forced exit of mitosis after prometaphase arrest with *S*-trityl-L-cysteine (63). Actin nucleation activity of centrosomes in these examples requires Arp2/3 complex. However, we found that FLARE formation was not affected by Arp2/3 inhibition. Thus, it is likely that FLARE is intrinsically different from these F-actin related structures around centrosomes in cultured human cells.

Some of fluorescently labeled phalloidin probes (7) and SiR-actin (65) can be used to monitor the fluorescence polarization of F-actin. Their live-cell applications are limited to specific biological preparations where these organic compounds can be delivered to cytoplasm (for example, isolated cells and monolayered culture cells). Fluorescently labeled phalloidin has been commonly used as an F-actin probe, but its strong F-actin stabilizing effect is not suitable for physiological studies in living cells where the activity of actin assembly/disassembly is essential. The successful application of POLArIS^act^ to study the dynamics of the actin cytoskeleton during early development of starfish eggs and embryos strongly suggests that POLArIS^act^ is a promising alternative to fluorescent phalloidin for F-actin studies in living cells.

Comparison to other widely-used F-actin probes proved that POLArIS^act^ is an excellent F-actin probe. In fixed cells, the localization of POLArIS^act^ was indistinguishable from that of phalloidin staining, which is currently thought to be the most reliable marker for F-actin. When observed in living cells, the cytoplasmic background signal from POLArIS^act^ was apparently lower than that of Lifeact or F-tractin and as low as that of UtrCH. We recently reported UtrCH-based F-actin probes for FPM, namely UG3 and UG7, can be used for orientation monitoring of F-actin with FPM (9). Similarly to POLArIS^act^, UG3 and UG7 are genetically encoded and the orientations of their fluorescence polarization are both parallel to the actin filament, but their polarization factor in living cells (<0.1) (9) was substantially lower than that of POLArIS^act^ (∼0.3, Fig. S2C). We expressed UG3 in starfish oocytes and found that the fluorescence polarization on the actin meshwork upon GV breakdown was substantially weaker than that of POLArIS^act^ (Fig. S9A). In addition, it was very difficult to recognize FLARE with FPM during cleavages by using UG3 (Fig. S9B and C), probably due to UG3’s low fluorescence polarization. Together, these results demonstrate that at present, POLArIS^act^ is the best tool for monitoring F-actin dynamics with FPM.

Since polarization is one of the most fundamental properties of light, POLArIS is expected to open a new avenue to various fluorescent spectroscopic applications such as, but not limited to, high throughput screening (66), super-resolution (13–16), resonance energy transfer (67, 68), and fluorescence correlation (69, 70). By taking advantage of the genetically encoded nature, POLArIS has the potential of supporting broader applications of fluorescence polarization techniques to a variety of cell types, species, and many types of specimens such as developing embryos and whole bodies of animals. Furthermore, POLArIS can be expressed in a cell-type/tissue specific manner when combined with specific promoters.

## Materials and Methods

### Cell culture

HeLa cells (a generous gift from Dr. Yoshimori, Osaka University) and HeLa M cells (a generous gift from Dr. Lowe, University of Manchester) were incubated at 37°C in Dulbecco’s modified Eagle medium (DMEM, nacalai tesque) containing 10% fetal bovine serum (FBS) with 5% CO_2_. LLC-PK1 cells (EC86121112-F0, ECACC) were incubated at 37°C in Medium 199 (Merck) containing 10% fetal bovine serum (FBS) with 5% CO_2_. XTC cells (a generous gift from Dr. Mitchell, Northwestern University) were incubated at room temperature in 70% Leibovitz’s L-15 medium (Thermo Fisher Scientific) containing 10% FBS.

### Plasmid construction

Sequence information of binding domains of Adhirons used in this study was described at the 60th Annual Meeting of Biophysics Society (21). DNA fragments of Ad- A, B, and C were synthesized by GeneArt Strings DNA Fragments (Thermo Fisher Scientific). The sequences for a fusion protein consisting of circularly permutated GFP (cpGFP; developed in our lab (9)) and each Adhiron were amplified by PCR using PrimeSTAR MAX (TAKARA Bio) and inserted in modified pcDNA3 vector (71), so that these two proteins are connected by a flexible linker. Then the flexible linker was replaced with 11 different linker sequences (L1-L11) using typical restriction enzyme cloning methods.

To make plasmids for mRNA synthesis, the POLArIS^act^ sequence and EMTB-mCherry (Addgene plasmid 26742) sequence were amplified by PCR and subcloned into the pGEMHE vector using Gibson Assembly (72)

### Protein purification and electrophoresis

The coding sequence for cpGFP, Ad-A, or POLArIS^act^ was inserted into the C-terminus of His tag in the pRSET B bacterial expression vector. *E. coli* JM109 (DE3) transformed with these plasmids were precultured in LB medium for 6 h at 37°C, diluted in LB medium at 1:1000, and cultured again for 72 h at 20°C. Cells of *E. coli* were collected by centrifugation and resuspended in TN buffer (50 mM Tris-HCl, 300 mM NaCl, pH 7.4). After cell disruption by sonication, the protein was purified using affinity purification with Ni Sepharose 6 Fast Flow (Cytiva). Samples were subjected to SDS-PAGE and the gels were stained with CBB staining using Quick-CBB PLUS (FUJIFILM Wako Pure Chemical).

### Crystallization and structure determination

Sequence coding POLArIS^act^ T57S mutant was cloned into pET24a bacterial expression vector, so that the N-terminal T7 peptide sequence was removed and the TEV cleavage site was inserted between the C-terminus of Adhiron and the N-terminus of His tag. The generated plasmid was transformed into *E.coli* Rosetta (DE3) and the protein was over-expressed overnight at 20°C in LB medium with induction by 0.3 mM IPTG. The cells were collected and were lysed using a sonicator. After centrifugation, the supernatant was applied on a HisTrap HP column (Cytiva) and the target protein was eluted with a linear gradient of imidazole. The His tag was then cleaved by TEV protease and was removed by a HisTrap HP column. The protein was further purified by ion-exchange on a HiTrap Q column (Cytiva) and by size-exclusion chromatography on a HiLoad 16/60 Superdex 75 pg column (Cytiva) in a buffer containing 25 mM Hepes-NaOH (pH 7.5), and 150 mM NaCl.

Crystals were grown at 20°C by the method of sitting drop vapor diffusion with a reservoir solution containing 0.1 M sodium acetate (pH 4.5-5.0) and 6-8% (v/v) PEG 4000. Crystals were cryo-protected with the reservoir solution containing 25% glycerol. Diffraction data were collected at a wavelength of 1.0 Å at SLS beamline X06DA (Villigen). Data were processed with the XDS programs (73) and the structure was solved by molecular replacement using the Phaser program (74) from the PHENIX programs (75), with the superfolder GFP (PDB ID:4LQT) and the Adhiron (PDB ID:4N6U) as the search models. The structural model was built into the electron density map using COOT (76) and refined using the PHENIX program and Refmac5 from the CCP4 program suite (77). The final statistics are summarized in Table S1.

### Structural modeling of cpGFP-Adhiron fusion

Modeling of the fusion of proteins was performed on UCSF Chimera (25). The structure of cpGFP was made by modifying the structure of the superfolder GFP (PDB ID:4LQT) and fused with Adhiron (PDB ID:4N6U) with manually constructed linker structures.

### Pull-down assay

G- and F-actin were prepared from rabbit skeletal muscle actin (AKL99, Cytoskeleton) following the manufacturer’s protocol. To depolymerize actin oligomers, actin protein was suspended to 0.4 mg/ml in general actin buffer (5 mM Tris-HCl, 0.2 mM CaCl_2_, 0.2 mM ATP, 0.5 mM DTT, pH 8.0) and incubated for 1 h on ice. After centrifugation at 4°C for 1 h, at 55,000 rpm using a fixed angle rotor S120AT2 (himac), the supernatant was used as a G-actin solution. To prepare F-actin solution, G-actin suspended in polymerization buffer (5 mM Tris-HCl, 50 mM KCl, 2 mM MgCl_2_, 0.2 mM CaCl_2_, 1.2 mM ATP, 0.5 mM DTT, pH 8.0) was incubated at room temperature for 1 h.

Proteins were mixed at the concentrations indicated in Fig. S4B in 200 µl of general actin buffer (for samples containing G-actin) or polymerization buffer (for samples containing F-actin). After incubation at room temperature for 1 h, 20 µl was taken from each sample as an “input” fraction, and the remaining 180 µl was added to 10 µl of washed beads of Ni Sepharose 6 Fast Flow and incubated at room temperature for 1 h with gently mixing. After centrifugation, the supernatant was collected as an “unbound” fraction. Precipitated beads were washed with general actin buffer (G-actin samples) or polymerization buffer (F-actin samples), followed by centrifugation again, and the supernatant was removed. The beads remained were used as “bound” fraction. Each fraction was mixed with SDS sample buffer, incubated at room temperature for 1 h, and then boiled at 95°C for 10 min. These samples were subjected to SDS-PAGE with 8% or 5-11% Bullet PAGE One Precast Gel (nacalai tesque) and the gels were stained using Quick-CBB PLUS.

Scanned images of gels from three independent sets of assays were quantified using ImageJ. *P* values were manually calculated on Microsoft Excel by applying Šidák multiple comparison correction to Student’s t-test, and difference with p < 0.01 was considered significant.

### Co-sedimentation assay

F-actin was prepared in the same way as in pull-down assays. F-actin and POLArIS^act^ were mixed in 50 µl of polymerization buffer with a concentration of 2.5 µM and 1-10 µM, respectively. After incubation at room temperature for 30 min, the mixture was centrifuged at 4°C for 1 h, at 55,000 rpm. The supernatant and the pellet were mixed with SDS sample buffer and were boiled at 95°C for 10 min. Ad-A and sfGFP instead of POLArIS^act^ were assayed in the same way as controls. These samples were analyzed by SDS-PAGE using home-made 8% or 12% acrylamide gel and the gels were stained using Quick-CBB PLUS.

Scanned images of gels from three independent sets of assays were quantified using ImageJ. Based on experimental assumptions that POLArIS^act^ binds to F-actin with 1:1 stoichiometry, which is determined from the result that approximately up to 2.5 µM POLArIS^act^ was co-sedimented with 2.5 µM F-actin, the dissociation constant (K_d_) was obtained according to the definition. Each actual value when the concentration of POLArIS^act^ was 1.5-2.5 µM in three sets of assays was substituted into the equation of definition to obtain K_d_,, and the average of values from three experiments was calculated.

### Plasmid introduction into cultured cells

cpGFP-Adhiron expression plasmids were transfected into HeLa or XTC cells cultured on glass-bottom dishes or coverslips using Lipofectamine LTX or 3000 (Thermo Fisher Scientific). The medium was replaced with fresh medium approximately 5 h after transfection. For HeLa M transfection, Polyethylenimine “Max” (24765-1, Polysciences) was used for transfection. For LLC-PK1, the NEPA21 electroporator (Nepagene) was used to introduce plasmids into cells. For live-cell imaging, the medium was replaced with DMEM/Ham’s F-12 without phenol red (nacalai tesque) or Medium 199 without phenol red (Thermo Fisher Scientific) containing 10% FBS before observation.

### Fixation and staining of cultured cells

To examine the co-localization of cpGFP-Adhiron and F-actin using fluorescently labeled phalloidin, transfected cells were washed with phosphate-buffered saline (PBS), and then fixed with 4% paraformaldehyde in PBS (pH 7.4). After permeabilization with 0.1% TritonX-100 for 5 min, the specimens were incubated with 0.5 µg/ml of phalloidin-Atto565 (94072, Merck), 2 µg/ml of Hoechst 33342 (04915-81, nacalai tesque), and 5% skimmed milk in PBS for 1 h at room temperature. Stained cells were washed with PBS and mounted in a mounting solution made of Mowiol 4-88 (ITW Reagents).

### *In vitro* preparation of F-actin and labeling

Purified human platelet actin (APHL99, Cytoskeleton) was polymerized following the manufacturer’s protocol and attached to polylysine coated coverslips by incubation with 2 µg/ml actin in actin polymerization solution. Then, specimens were incubated with 10 nM purified recombinant POLArIS^act^ protein or 2-5 nM Alexa Fluor 488 Phalloidin (AF488-phalloidin, A12379, Thermo Fisher Scientific) and 10 mg/ml bovine serum albumin (BSA) in actin polymerization solution for 5 min.

### Starfish egg preparation and drug treatment

*Asterina pectinifera* starfishes were collected on the Pacific coast of Japan and were kept in laboratory aquaria with seawater at 14–15°C. Immature oocytes of *Asterina pectinifera* starfishes were treated with cold calcium-free seawater to remove follicle cells, and then incubated in filtered seawater. Oocytes were microinjected with the mRNA encoding POLArIS^act^, UG3, and/or EMTB-mCherry (synthesized *in vitro* from linearized DNA templates using the mMessage mMachine T7 ULTRA Transcription kit (Thermo Fisher Scientific)) on the day before observation and incubated in filtered or artificial seawater (ASW: 423 mM NaCl, 9 mM KCl, 23 mM MgCl_2_, 25 mM MgSO_4_, 9 mM CaCl_2_, 10 mM HEPES, pH 7.4) at 16°C overnight. After POLArIS^act^ and/or EMTB-mCherry were expressed at the appropriate level, oocytes were treated with 1-Methyladenine (1-MA, KANTO CHEMICAL) to induce maturation.

For observation of early development processes, oocytes were inseminated after germinal vesicle breakdown (GVBD) and incubated at 16°C until microscopic imaging was started. In pharmacological experiments, embryos were treated with 20 µM nocodazole (NCZ) (487929, Merck), 10 µM cytochalasin D (CytoD) (C8273, Merck), 200 µM CK666 (182515, Merck) or 10 µM SMIFH2 (344092, Merck) in ASW at the indicated timing (see figure legends for details), or perfused with 10 µM SMIFH2 dissolved in ASW by inserting a microneedle into the perivitelline space after fertilization envelope formation. In control experiments, embryos were exposed with ASW containing DMSO. Drugs were diluted in ASW from stock solutions dissolved in DMSO (NCZ, 10 mM; CytoD, 10 mM; CK666, 100 mM; SMIFH2, 50 mM) and the same volume of DMSO as the volume of each stock solution was added to ASW as control.

For observation of fertilization, 1-MA treated eggs were inseminated after GVBD. To perform simultaneous observation of F-actin and DNA, immature eggs were pre-stained with 10 µg/ml Hoechst 33342 for 30 min. Stained eggs were washed with ASW and then treated with 1-MA. Sperms were also stained with 10 µg/ml Hoechst 33342 for 5 min just before insemination.

### Fixation and fluorescent phalloidin staining of starfish embryos with PFA or PFA+GA

As soon as the green/magenta cross pattern clearly appeared, starfish embryos expressing POLArIS^act^ were fixed for 1 h with 4% PFA in MES buffer (10 mM MES, 150 mM NaCl, 5 mM EGTA, 5 mM glucose, 5 mM MgCl_2_, pH 6.1) or 0.25% GA and 4% PFA in HEPES buffer (10 mM HEPES, 460 mM NaCl, 10 mM KCl, 36 mM MgCl_2_, 17 mM MgSO_4_, pH 8.2) under the FPM. For AF488-phalloidin staining, embryos fixed with GA+PFA in HEPES buffer were then washed with HEPES buffer, and PFA and GA were quenched with 100 mM glycine. After permeabilization for 15 min with HEPES buffer containing 0.1% TritonX-100, embryos were stained for 2 h with AF488-phalloidin.

### Immunofluorescence of fixed starfish embryos

Starfish embryos were fixed for 1 h with a 3% v/v glyoxal (17226-95, nacalai tesque) fixative solution prepared as described previously (35). After fixation, embryos were washed with PBSw/NaCl (same as PBS except that NaCl concentration was adjusted at 460 mM) and glyoxal was quenched with 100 mM glycine. Embryos were then permeabilized with PBSw/NaCl containing 0.1% TritonX-100, blocked with 0.05% BSA, and sequentially incubated with primary and secondary antibodies dissolved in PBSw/NaCl. To visualize microtubules, YL1/2 rat anti-α-tubulin (MCA77G, Bio-Rad) was used as a primary antibody, and goat anti-rat IgG conjugated with Alexa488 (A-11006, Thermo Fisher Scientific) was used as a secondary antibody. To visualize DNA and F-actin, Hoechst 33342 and phalloidin-Atto565 were used, respectively. Stained embryos were placed on a MAS-coated slide glass (Matsunami Glass), mounted with SlowFade Diamond (S36963, Thermo Fisher Scientific), covered with a coverslip, and sealed with nail polish. The observation was carried out within 2 days, as staining with phalloidin-Atto565 was blurred 3 days after glyoxal fixation.

### Fluorescence microscopy

Fluorescence microscopic observation of fixed HeLa cells was performed with the Nikon Eclipse E600 microscope with a dry objective lens (PlanFluor 40x 0.75 N.A., Nikon) or an oil immersion objective lens (PlanFluor, 100x 1.30 N.A., Nikon). Images were acquired with the application software ACT-2U (Nikon), and then processed with ImageJ.

Fluorescence microscopic observation of living starfish eggs/embryos and data acquisition were performed with Nikon Eclipse TiE microscope with dry objective lenses (PlanApo 10x 0.45 N.A., and 20x 0.75 N.A., Nikon) and silicone oil immersion objective lenses (UPLSAPO, 30x 1.05 N.A., and 40x 1.25 N.A., Olympus) in a room kept at 20°C.

Fluorescence microscopic observation of fixed starfish eggs/embryos and data acquisition were performed with Nikon C1 CLSM with a silicone oil immersion objective lens (UPLSAPO, 40x 1.25 N.A., Olympus), Leica SP8 CLSM (Leica) with an oil immersion objective lens (HC PL APO 63x/1.40 Oil CS2, Leica), or Carl Zeiss LSM510 CLSM with a water immersion objective lens (C-Apochromat 40x/1.2 W Corr, Carl Zeiss). Images were taken with NIS-Elements Advanced Research, Leica LAS X, and Zeiss ZEN, respectively, and processed with ImageJ.

### Two-axis Fluorescence Polarization Microscopy

The polarization beam splitting system was assembled at the detection port of Nikon Eclipse TiE (9). In brief, samples were illuminated by the isotropically polarized LED light from SPECTRA light engine (lumencor) and the linear polarization beam splitter splits fluorescence into two orthogonal polarization orientations, 0° (horizontal orientation) and 90° (vertical orientation), and detected with a single camera (iXon3 897 EM-CCD (Andor Technology) or Zyla 4.2 plus sCMOS (Andor Techonology)). Both polarization images were labeled with pseudo-colors (horizontal polarization, green; vertical polarization, magenta) with NIS-Elements Advanced Research. Images of merged color were shown in real-time with the dual-view function of NIS-Elements Advanced Research. We used dry objective lenses (PlanApo 10x 0.45 N.A., and 20x 0.75 N.A., Nikon) and silicone oil immersion objective lenses (UPLSAPO, 30x 1.05 N.A., and 40x 1.25 N.A., Olympus). iXon3 897 EM-CCD camera (Andor Technology) and Zyla 4.2 plus sCMOS (Andor Techonology) were used for detection. Filter cube LF488-C (Semrock) was used for cpGFP-Adhiron fluorescence.

### Instantaneous FluoPolScope

Imaging and analysis using Instantaneous FluoPolScope were performed as reported previously (7, 9). The excitation illumination achieves isotropic polarization at the specimen plane. Fluorescence images were collected with a high NA objective lens (PlanApo TIRF 100x 1.49NA oil, Nikon). In the collimated space, linear polarization beam splitters separate the fluorescence into four images and analyze their linear polarization along 0°, 45°, 90°, and 135° orientations. Based on the intensity imbalance of polarized fluorescence along four orientations, polarization factor and the orientation of maximum polarization of the objects are computed.

### Image analysis for fluorescent microscopy

The intensity profiles of fixed HeLa cells were obtained using ImageJ.

The length of actin bundles (from the point on the cell surface where the bundle appears to the tip of the bundle) in starfish oocytes during fertilization were measured using the Nikon imaging software NIS-Elements Advanced Research. Analyses were performed on 8 different zygotes.

### Calculation of the polarization value (PV)

For quantification of fluorescence anisotropy of cpGFP-Adhiron constructs, transfected HeLa M cells were arrested in mitosis using the thymidine-NCZ method. In brief, HeLa M cells were treated with 2 mM thymidine for 20 h, and released for overnight in the presence of 100 ng/mL NCZ. We defined Polarization Value (PV) as

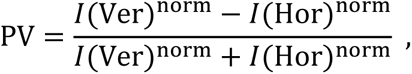

where *I*(Ver)^norm^ = [cortex ROI intensity (Ver) – background (Ver)], *I*(Hor)^norm^ = [cortex ROI intensity (Hor) – background (Hor)]·CF, and CF = total intensity (Ver) / total intensity (Hor). The mean intensity of a small rectangle ROI enclosing vertical cortex in the cross-section of a mitotic cell was measured both in vertical polarization and horizontal polarization images to calculate cortex ROI intensity (Ver) and cortex ROI intensity (Hor), respectively. The mean intensity of a square ROI outside of the cell was measured both in the same vertical polarization and horizontal polarization images to calculate background (Ver) and background (Hor), respectively. CF (correction factor) was obtained by calculating the ratio of the mean ROI intensities between vertical and horizontal polarization images that contain a whole cell, using images of cells expressing cpGFP-Ad-A flex. Examples of ROI settings are shown in Fig. S2A. Positive PV values indicate the polarization is parallel to the axis of actin filaments, and negative PV values indicate the perpendicular polarization. n = 10 for each calculation. *P* values against cpGFP-Ad-A (flex) were manually calculated on Microsoft Excel by applying Šidák multiple comparison correction to Student’s t-test. Difference with p < 0.01 was considered significant.

### Analysis of the cross pattern of FLARE

The presence of the green (horizontal)/magenta (vertical) cross patterns and their centers were deduced using the strategy as detailed below. Suppose a polar coordinate system (r, φ) is set over the radial F-actin array, and the transition dipole moments (TDMs) of the fluorophores of POLArIS^act^ are aligned on each F-actin filament. When the fluorophores are excited by vertical polarization (parallel to line φ = 0), the fluorescence efficiency of each fluorophore varies with the angle between the polarized excitation and the TDMs. Thus, the distribution of observed fluorescence intensity Fl_vertical_ can be modeled as:

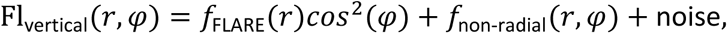

where *f*_FLARE_(*r*) is the radial profile of the fluorescent component that is aligned on radial F-actin array, and *f*_non-radial_(*r*, *φ*) is the fluorescent component that is not radially aligned. The distribution of fluorescence intensity Fl_horizontal_ to be observed with horizontal excitation will be:

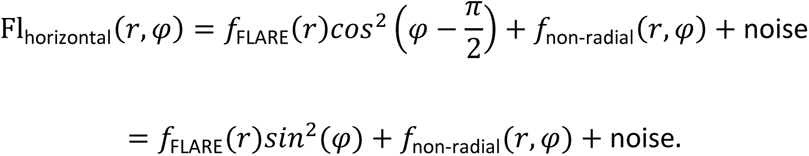

Then the polarization differential image intensity Fl_diff_ of these two will be modeled as:

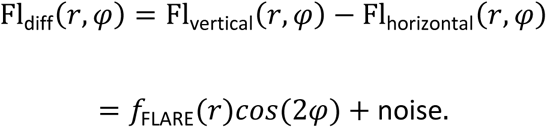

Thus, fluorophores aligned on a radial F-actin array show a fluorescence intensity pattern that varies with cos(2φ) when measured along the circle of fixed radius around the center of the array in the differential image. Conversely, if this characteristic pattern is found on the polarization differential image, it suggests the possible presence of a radial F-actin array there.

Based on this, we used the following algorithm to search for the presence of cross patterns. First, we created a kernel image with this characteristic intensity pattern of cos(2φ):

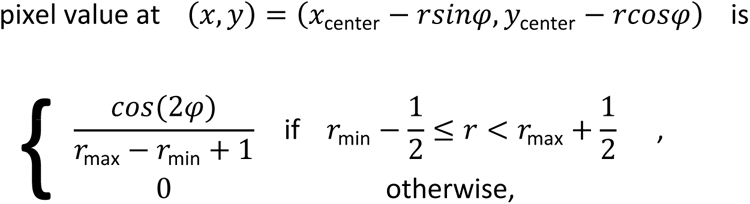

where (*x*_center_, *y*_center_) is the center of the kernel image, and *r*_min_ or *r*_max_ specifies the size of the concentric area to look for. Then, we convolved the polarization differential images with this cos(2φ) kernel image. Each pixel value of the convolved result is related to the likelihood of the presence of the cos(2φ) pattern with the center at that pixel. We inspected several points with local maximal values as candidates and determined the center of radial F-actin arrays.

The radial extent of the F-actin array was not readily discernable because it was blurred in the noisy background. So, to help visualize the radial profile, we rotated the polarization differential image around the center of the array and calculated the autocorrelation at each radius *G*_rot_:

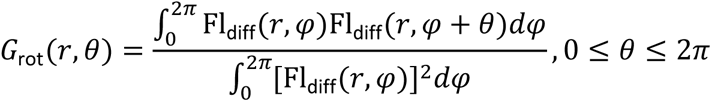

The rotational autocorrelation above was normalized to have a value of 1 at *θ* = 0 or *θ* = 2*π*. Bicubic interpolation was used for subpixel sampling when calculating the autocorrelation of pixel values from the differential image. The rotational autocorrelation of the radial FLARE component becomes:

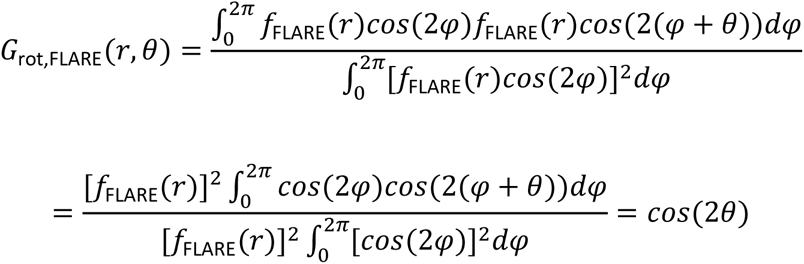

This curve takes a local maximum at *θ* = *π* and local minima at 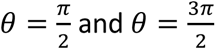. In contrast, the autocorrelation of the noise component is expected to have a sagging profile with a minimum at *θ* = *π*. The shape of *G*_rot_(*r*, *θ*) surface calculated from actual differential images will depend on the relative intensity of the radial FLARE component and the noise, but the presence of a peak at *θ* = *π* in the rotational autocorrelation will be a good indicator of the presence of the cross pattern at the radius *r*. The algorithms described above were packaged as an ImageJ plugin (S4SSH) and deposition of the plugin at GitLab is in progress.

Obtained 2D-plots of the rotational autocorrelation for each radius were converted to 3D-plots using ImageJ’s ‘surface plot’ function.

## Acknowledgments

We are grateful to N. Nakai (TMDU) and H. T. Miyazaki (NIMS) for helpful discussion, O. Hoshi (TMDU) for continuous encouragement, and to M. Taguchi (TMDU) for secretarial and experimental assistance. We deeply thank H. Iwasaki (Gunma-U) for suggestion making the initial opportunity of the collaborative research using starfish oocytes, R. Nitta and M. Aoki (RIKEN) for discussion and initial trial for the crystallization of POLArIS^act^, K. Hanada and M. Inoue (RIKEN) for plasmid preparation for the expression of POLArIS^act^ protein in bacterial cells, and T. M. Watanabe (RIKEN-QBiC) for discussion and initial assistance for FPM construction. We are grateful to the late Dr. Shinya Inoué (1921 - 2019), the father of polarized light microscopy, for his great support and his pioneering work of polarized fluorescence in green fluorescent protein crystals (78).

## Data Availability

All data are included in the manuscript and supporting information. The deposition of ImageJ plugin for FLARE analysis at GitLab is in progress.

## Author Contributions

Discussion with T.T. led S.T. to start a close collaboration with T.T. and to conceive of the project. A.S. and Kei.S. conducted most experiments and performed analysis with other collaborators. K.C. performed injections into starfish oocytes and supervised the related experiments. Kei.S. and Ken.S. jointly constructed the two-axis FPM along with the guidance by T.T. M.K. contributed imaging analysis for starfish embryo data. S.M. contributed instantaneous FluoPolScope analysis tools. Y.T. worked with N.S. to collect the crystallography data and determined structures under the supervision by M.S. T.T. conducted experiments and performed analysis by instantaneous FluoPolScope with assistance of H.I. S.T. designed experiments and supervised the project. A.S. and Kei.S. jointly wrote the manuscript with inputs from all authors.

## Supporting Information

**Fig. S1.**
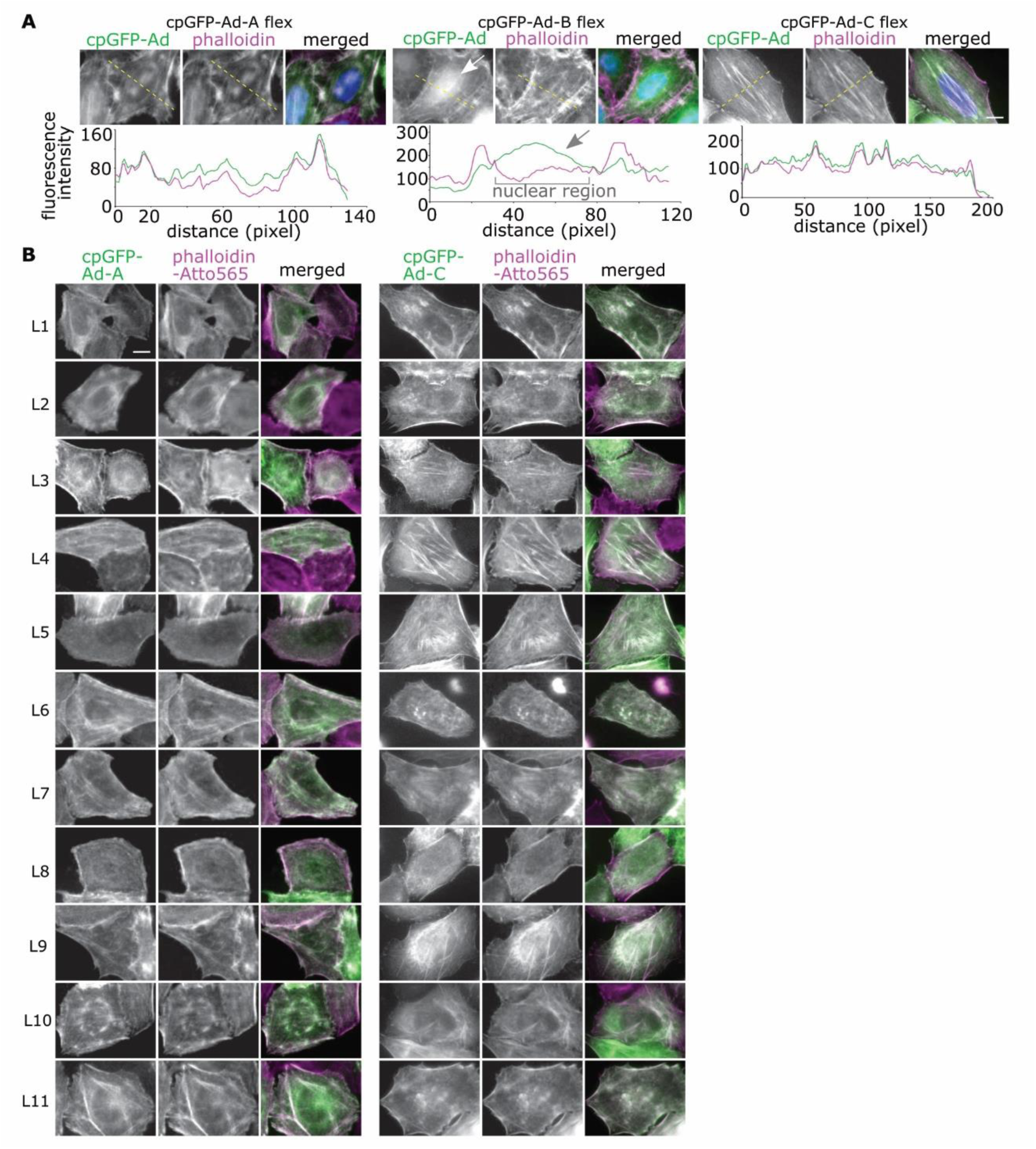
**Localization analysis of cpGFP-Adhiron constructs.** (A) Top row: Representative fluorescence microscopy images of fixed HeLa cells expressing cpGFP-Ad-A/B/C flex (left panels) with phalloidin-Atto565 (middle panels) staining. Merged images of cpGFP-Adhiron (green), phalloidin-Atto565 (magenta), and DNA stained with Hoechst (blue) are also shown in the right panels. Bottom row: Intensity profiles for cpGFP-Ad-A/B/C and phalloidin-Atto565 channels along the dashed yellow lines marked on the images in top panels. Arrows show a high background in the nuclear region of cpGFP-Ad-B flex fluorescence. Scale bar: 10 µm. (B) Co-localization analysis of cpGFP-Adhirons with actin filaments stained with phalloidin. Representative fluorescence microscopy images of fixed HeLa cells expressing cpGFP-Ad-A/C L1-L11 (left columns) with phalloidin-Atto565 (middle columns) staining are shown. Merged images of cpGFP-Adhiron (green) and phalloidin- Atto565 (magenta) are shown in the right columns. Scale bar: 10 µm.

**Fig. S2.**
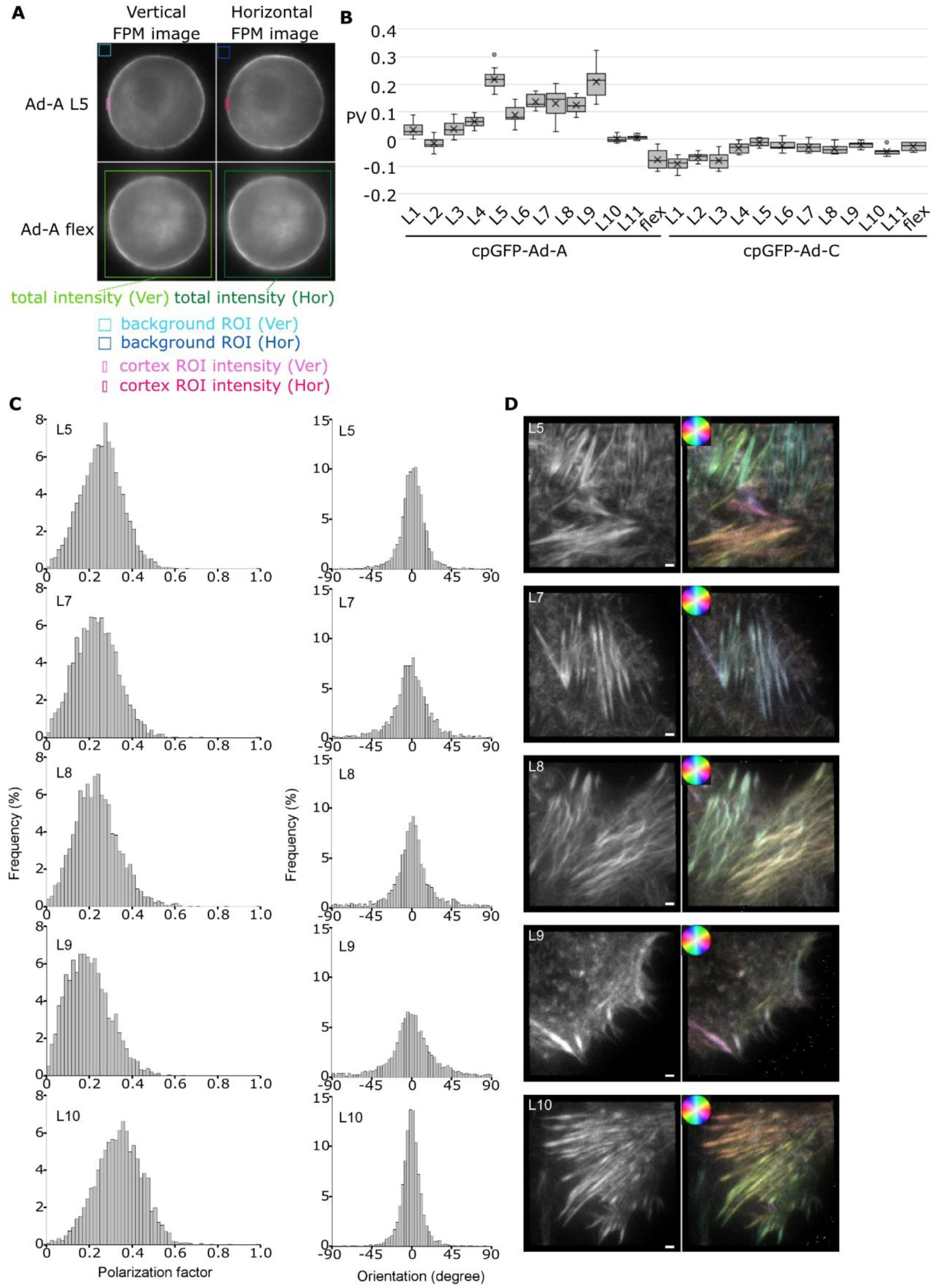
**Screening of cpGFP-Adhiron constructs by quantifying polarization of fluorescence in living cells.** (A) Examples of ROI settings in two-axis FPM images for calculation of Polarization Value (PV) of cpGFP-Ad-A L5 are shown. See Materials and Methods for details of PV calculation. (B) PVs for all cpGFP-Ad constructs are shown by boxplots with cross marks indicating averages. Statistics: Student’s t-test with Šidák correction. **, P < 0.01. n=10. (C) Instantaneous FluoPolScope analysis of five cpGFP-Ad-A constructs (L5, 7, 8, 9, and 10) expressed in LLC-PK1 cells. Polarization factor (left) and polarization orientation (right) are shown in histograms. The ordinates of histograms are the frequencies of detected particles. Orientation at 0° indicates the polarization orientation of the particle is parallel to actin filaments, and orientation at −90° or 90° indicates the polarization orientation of the particle is perpendicular to actin filaments. (D) Representative images of instantaneous FluoPolScope analysis. Left panels show the ensemble fluorescence intensity of four polarization orientations, and right panels show the orientation of ensemble polarization by pseudo-color. The hue color circles in the right panels show the relationship between pseudo-color and orientation of ensemble polarization. Scale bar: 1 µm.

**Fig. S3.**
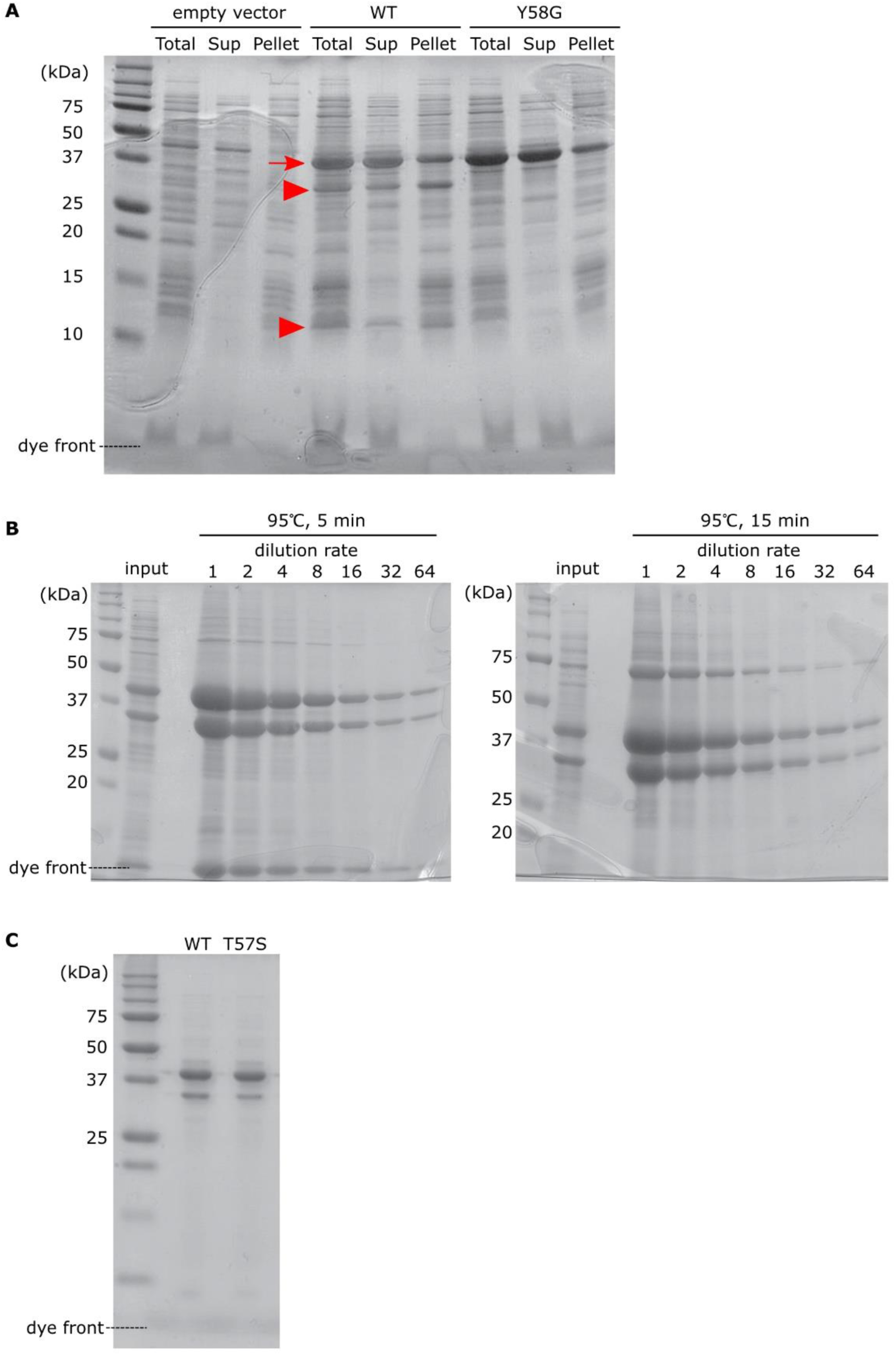
**Selection of POLArIS^act^ mutants suitable for structure determination.** (A) CBB-stained SDS-PAGE gel of whole protein (Total), supernatant (Sup), and pellet of *E. coli* JM109 (DE3) bacterial cells expressing POLArIS^act^ or its Y58G variant. POLArIS^act^ protein (WT) unexpectedly produced extra bands at 32 kDa and 11 kDa (arrowheads) in addition to the expected band of 40 kDa (arrow). Extra bands disappeared when the Y58G mutant, which is expected to prevent cpGFP from forming the chromophore, was used, indicating that extra bands were produced due to “backbone fragmentation,” during chromophore formation or maturation. (B) CBB-stained SDS-PAGE gels of purified POLArIS^act^. Proteins were purified from POLArIS^act^-expressing bacterial cells and prepared samples were heated at 95°C for 5 min (left) or 15 min (right) and loaded onto gels with sequential dilutions. The production of extra bands shown in (A) is unlikely to be caused by the incompletion of protein denaturation, as the intensities of extra bands were not affected by the difference in the length of the heating. (C) CBB-stained SDS-PAGE gel of purified wild-type POLArIS^act^ (WT) and its mutant (T57S) proteins. T57S mutant of POLArIS^act^ was comparatively resistant to “backbone fragmentation” and used for the crystal structure analysis.

**Fig. S4.**
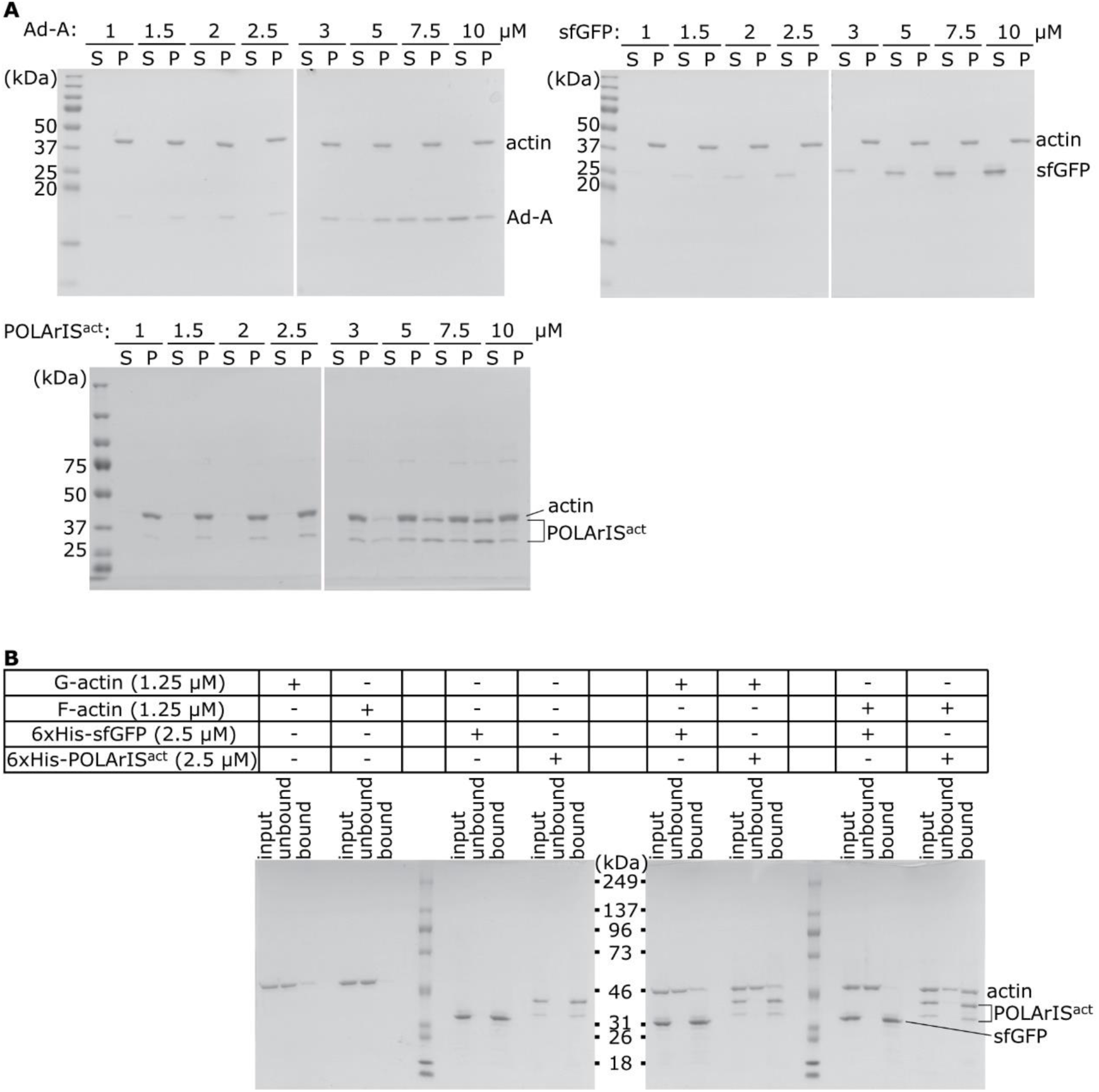
**Co-sedimentation and pull-down assays to analyze interaction between POLArIS^act^ and F-actin.** (A) CBB staining of typical SDS-PAGE gels of a co-sedimentation assay in which 2.5 µM F-actin were mixed with increasing concentrations of recombinant Ad-A (top left), sfGFP (top right) or POLArIS^act^ (bottom). Lanes show supernatant (S) and pellet (P) of each mixture with different concentrations of proteins. (B) CBB staining of typical SDS-PAGE gels of actin pull-down assay. Components of each reaction are shown in the list. The bound fraction was twice as concentrated as input and unbound fractions.

**Fig. S5.**
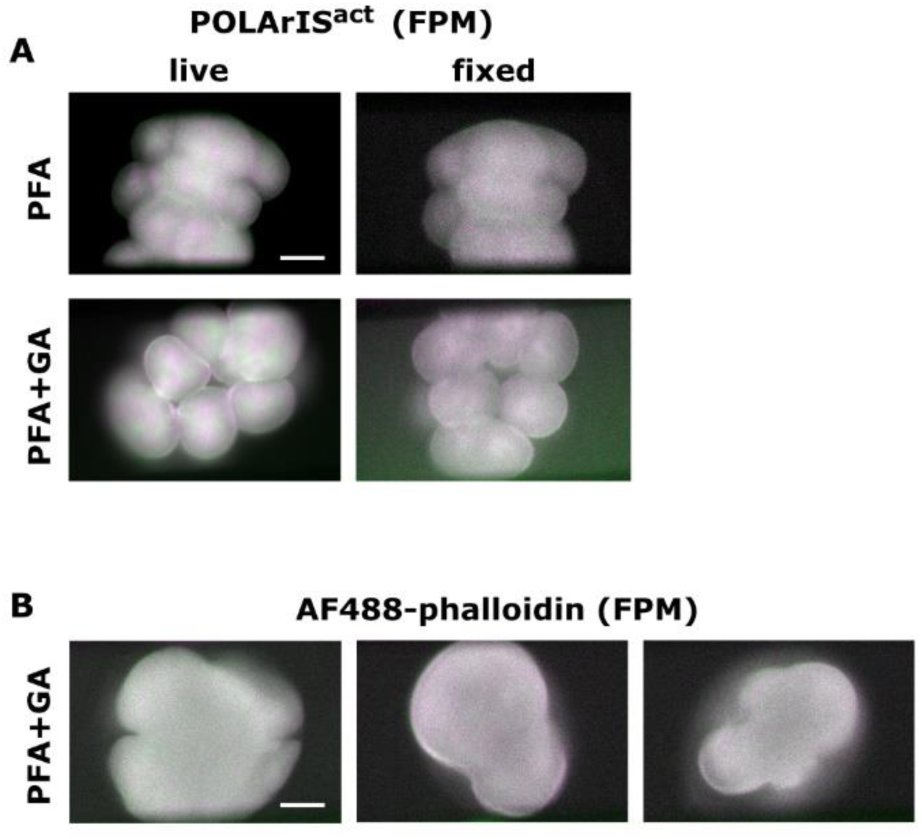
**Fluorescence polarization of FLARE was not preserved with PFA or PFA+GA fixatives.** (A) Green/magenta two-axis FPM images of live and fixed starfish embryos expressing POLArIS^act^ in the fourth cleavage. Soon after the green/magenta cross pattern appeared (live), the same embryos were fixed with PFA or PFA+GA (fixed) by perfusing fixative. The cross patterns of POLArIS^act^ were disappeared after fixation. (B) Three examples of green/magenta two-axis FPM images of AF488-phalloidin staining of PFA+GA-fixed starfish embryos expressing POLArIS^act^. The cross pattern was not visible in all embryos. Scale bar: 50 µm.

**Fig. S6.**
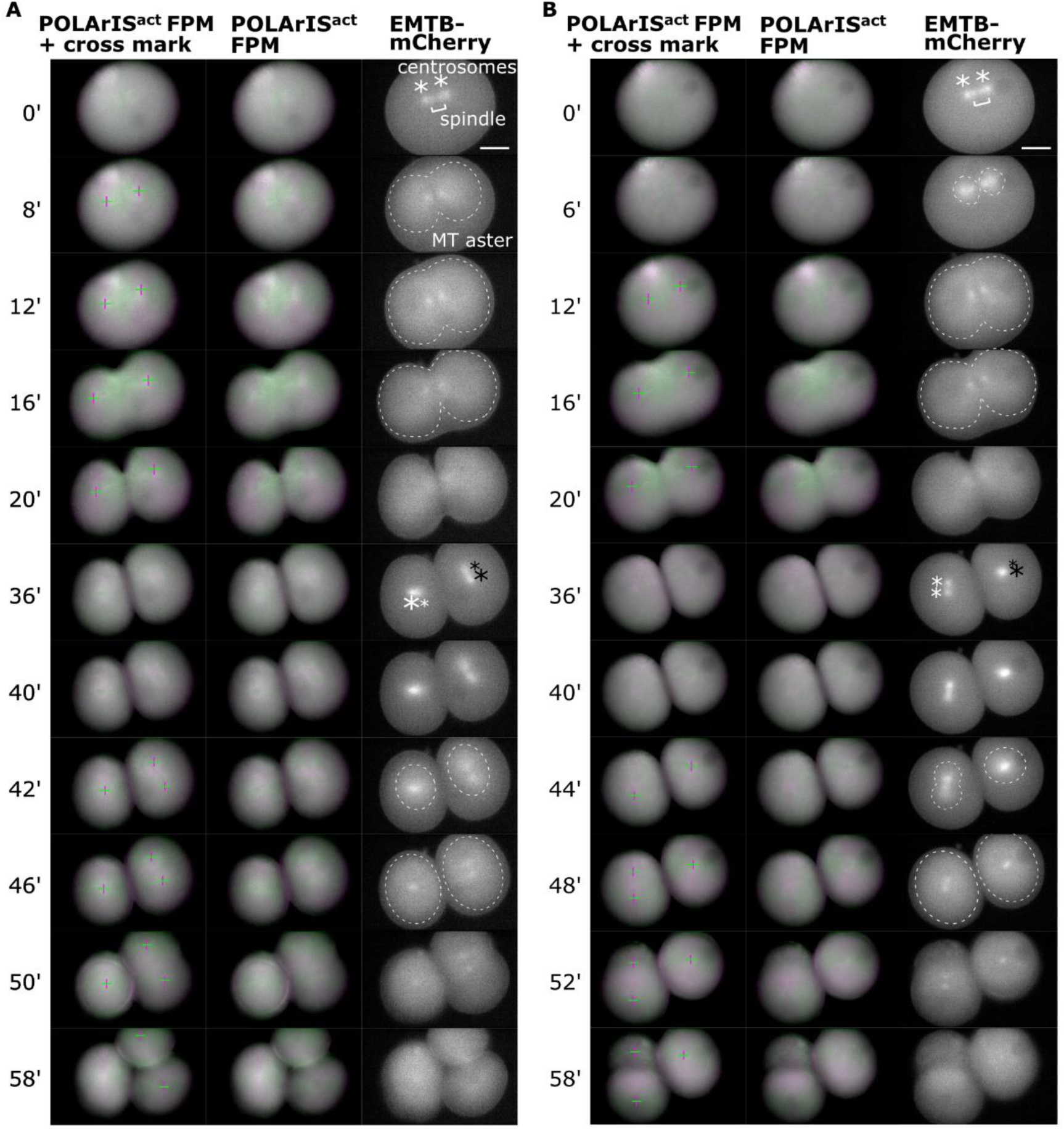
**Similar distribution and dynamics of FLARE and microtubule asters.** (A) Longer time-lapse green/magenta (left and middle columns) two-axis FPM images of the first and the second cleavages of the same starfish embryo expressing POLArIS^act^ and EMTB-mCherry as in Fig. 5C. The right column shows vertical polarization images of EMTB-mCherry. Cross marks (left column) indicate centers of green/magenta cross patterns. Centrosomes are shown by asterisks and contours of microtubule asters are shown by dashed lines. In POLArIS^act^ images, the background signal was subtracted for a better presentation of cross patterns. (B) Another example of time-lapse green/magenta (left and middle columns) two-axis FPM images of the first and the second cleavages of a starfish embryo expressing POLArIS^act^ and EMTB-mCherry. Scale bar: 50 µm.

**Fig. S7.**
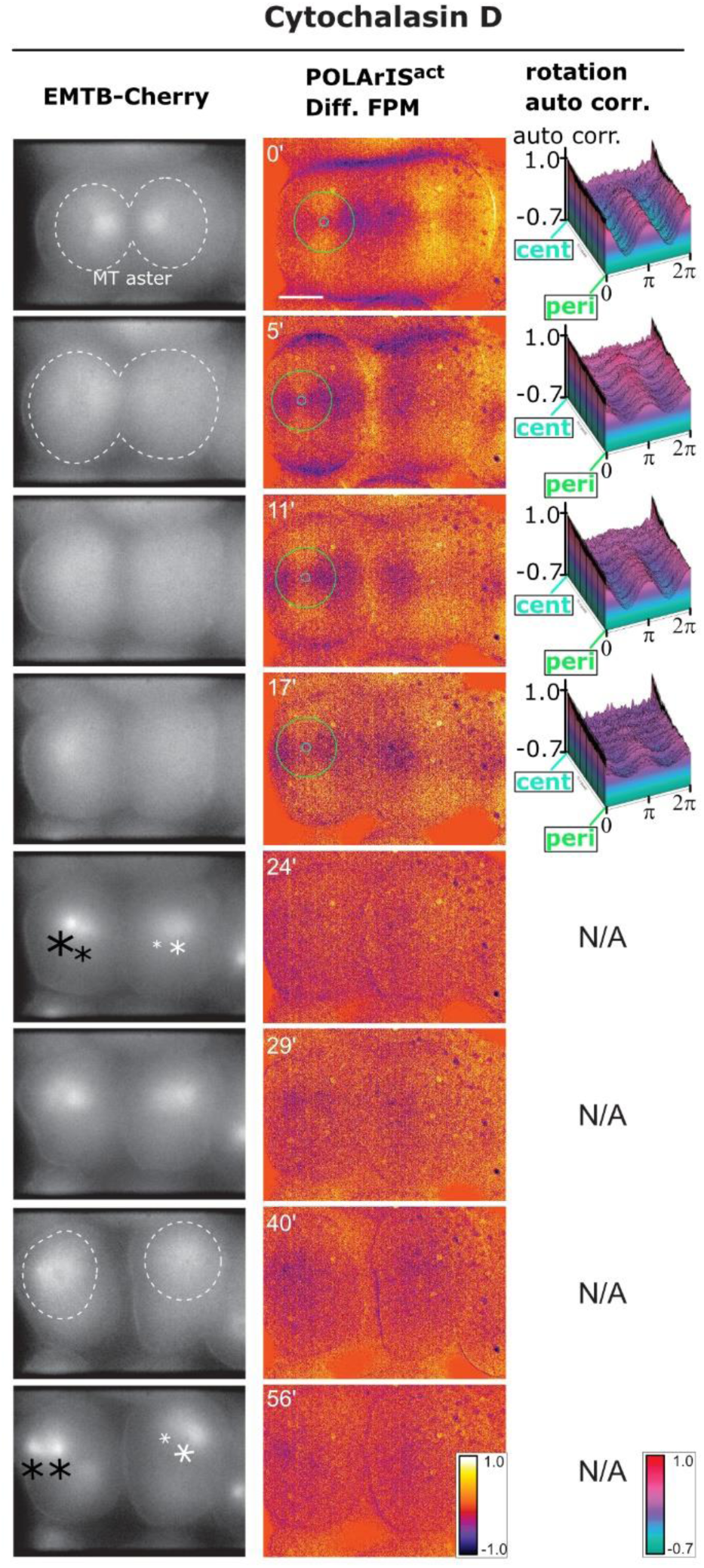
**Microtubule aster dynamics does not depend on FLARE.** Time-lapse two-axis FPM observation of a cytochalasin D-treated embryo expressing POLArIS^act^ and EMTB-mCherry in the fourth cleavage. The drug was added at time 0. The left column shows vertical polarization images of EMTB-mCherry, the middle column shows differential FPM images of POLArIS^act^ with circles of min/max diameters used in autocorrelation analyses, and the right column shows 3D-plots of autocorrelation analyses of FLARE. Asterisks indicate centrosomes, dashed lines indicate contours of microtubule asters. Scale bar: 50 µm.

**Fig. S8.**
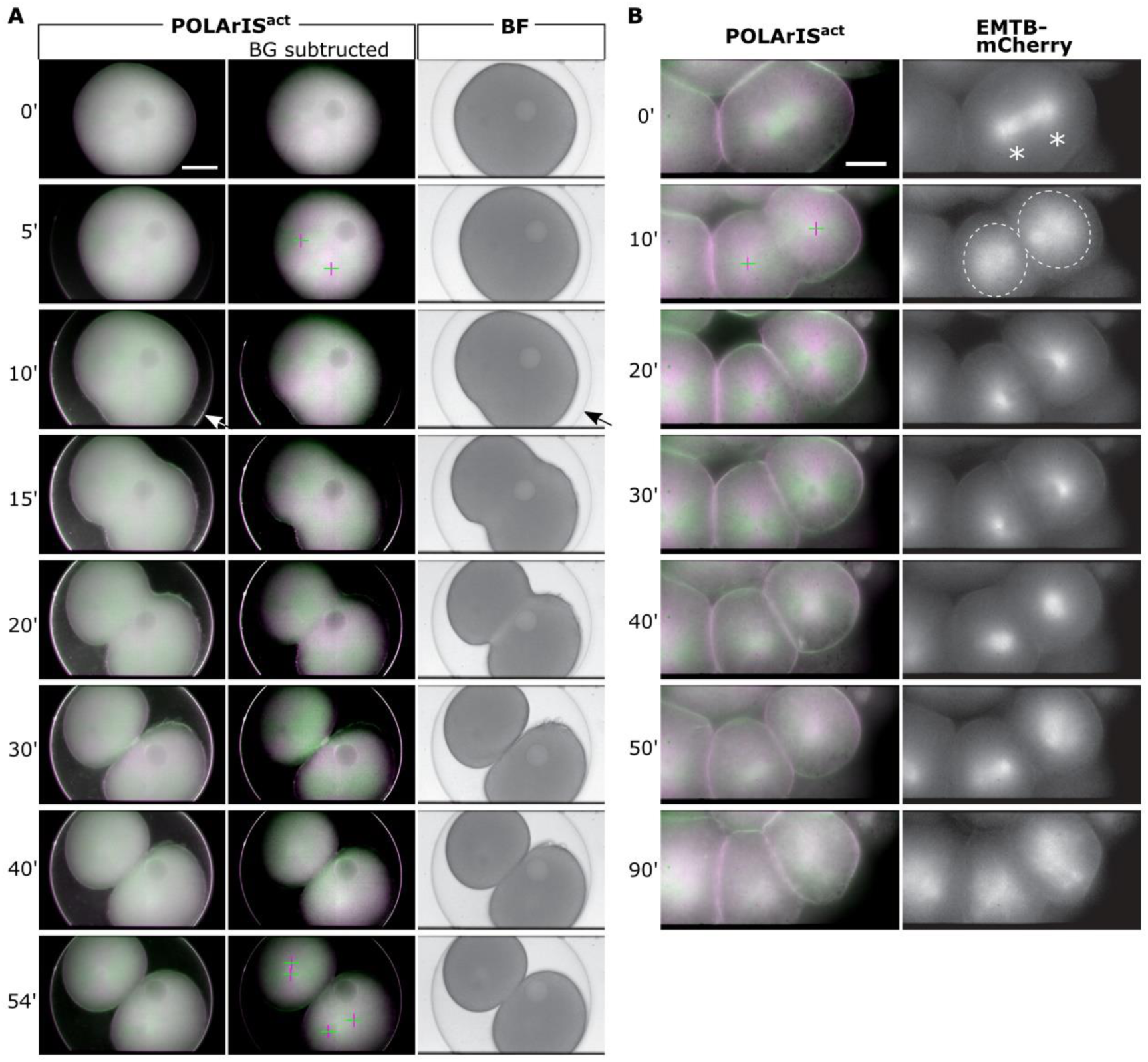
**Time lapse observations of SMIFH2 treated starfish embryos.** (A) A starfish embryo expressing POLArIS^act^ was treated with SMIFH2 before the first cleavage (time 0). Images by two-axis FPM (left and middle columns) and their bright-field view (right column). In panels of the middle column, the background signal was subtracted for the better presentation of green/magenta cross patterns. Arrows indicate the fertilization envelope and cross marks indicate the centers of the cross patterns of FLARE structures. Accumulation of the drug on the fertilization envelope was evident 10 min after the drug addition. Scale bar: 50 µm. (B) A starfish embryo expressing POLArIS^act^ and EMTB-mCherry was treated with SMIFH2 and observed with two-axis FPM. SMIFH2 was microinjected and perfused under the elevated fertilization envelope. The drug was added just after microtubule asters started to extend during the fourth cleavage (time 0). The left column shows green/magenta FPM images of POLArIS^act^ and the right column shows vertical polarization images of EMTB-mCherry. Cross marks indicate the centers of the green/magenta cross patterns. Asterisks indicate centrosomes, dashed lines indicate contours of microtubule asters. Scale bar: 25 µm.

**Fig. S9.**
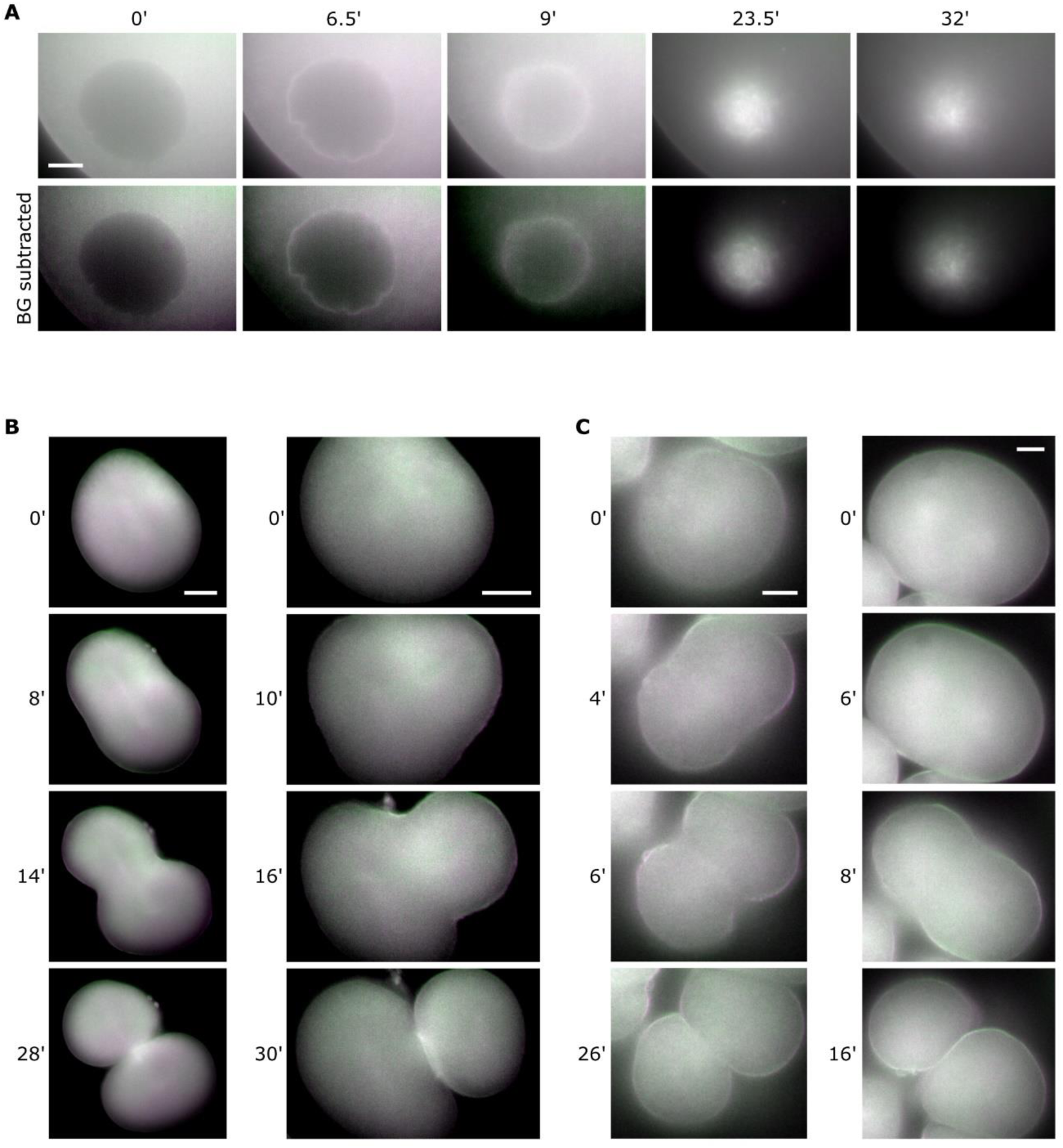
**Time lapse FPM observations of starfish eggs expressing UG3.** (A) Time-lapse green/magenta two-axis FPM images of starfish oocytes expressing UG3 during GV breakdown. Time 0 is just before the breakdown, approximately 30 min after 1-MA addition. In bottom panels, the background signal was subtracted for the better presentation of actin structures. (B, C) Time-lapse green/magenta two-axis FPM images of embryo expressing UG3 in the second (B) or the fourth cleavages (C). Scale bars: 25 µm (A, C), 50 µm (B).

**Table S1.** Data Collection and Refinement Statistics.

|  |  |
| --- | --- |
| ***Data collection and processing*** |  |
| Beamline | X06DA, SLS |
| Space group | *I* 2<sub>1</sub> 2<sub>1</sub> 2<sub>1</sub> |
| Unit-cell parameter |  |
| $a, b, c$ (Å) | 78.1, 113.6, 132.7 |
| $\alpha, \beta, \gamma$ (°) | 90.0, 90.0, 90.0 |
| Wavelength (Å) | 1.000 |
| Resolution range (Å) | 50.0–2.50 (2.64–2.50) |
| Redundancy | 7.1 (7.1) |
| Completeness (%) <sup>a</sup> | 99.9 (99.8) |
| $R_{\text{sym}}$ <sup>b</sup> (%) <sup>a</sup> | 1.62 (9.6) |
| $I/\sigma(I)$ <sup>a</sup> | 13.3 (1.0) |
| No. monomers/asymmetric unit | 1 |
| ***Model refinement*** |  |
| No. of reflections | 20785 |
| No. of protein atoms | 2528 |
| No. of water molecules | 81 |
| $R_{\text{work}}/R_{\text{free}}$ (%) | 21.4/23.8 |
| r.m.s.d. for bond length (Å) | 0.003 |
| r.m.s.d. for bond angles (°) | 0.633 |
| ***Residues in the Ramachandran plot*** |  |
| Favored region (%) | 95.7 |
| Allowed regions (%) | 2.6 |
| PDB ID | XXXX |
| <sup>a</sup> Statistics for the highest resolution shell are given in parentheses. |  |
| <sup>b</sup> $R_{\text{sym}} = \sum_{hkl} \sum_i \vert I_i(hkl) - \langle I(hkl) \rangle\vert / \sum_{hkl} \sum_i I_i(hkl)$. | |

**Movie S1.**

A representative movie of the single-particle observation of POLArIS^act^ with Instantaneous FluoPolScope in a lamellipodium of a living XTC cell expressing POLArIS^act^. Left panels show the ensemble fluorescence intensity of instantaneous FluoPolScope and right panels show the polarization factor and the orientations of polarized fluorescence (indicated by yellow lines, yellow scale bar shows the maximum polarization factor) of detected POLArIS^act^ particles (yellow circles). x312 actual speed.

**Movie S2.**

Time-lapse of two-axis FPM (green/magenta) of starfish oocytes expressing POLArIS^act^ during the breakdown of the GV and formation of the F-actin meshwork, related to Fig. 3A. Time 0 is just before the breakdown, about 27 min after 1-MA addition. Arrow indicates GV, and arrowheads indicate actin filaments with clear polarization in the meshwork (green: horizontal, magenta: vertical). x236 actual speed.

**Movie S3.**

Time-lapse of starfish oocytes expressing POLArIS^act^ during the polar body extrusion with two-axis green/magenta FPM (left) and bright-field view (right), related to Fig. 3B. The positions of the polar body extrusion and the accumulation of contractile ring-like F-actin are shown by arrowhead and arrow, respectively. x312 actual speed.

**Movie S4.**

Fluorescence microscopy time-lapse of the fertilization of starfish oocyte expressing POLArIS^act^ with Hoechst staining, related to Fig. 3D. Top shows the entire egg, and bottom magnifies the area where the actin bundle, DNA of the egg (arrowheads) and sperm (arrows) were observed. Left: POLArIS^act^, middle: Hoechst, right: merged (POLArIS^act^: green, Hoechst: magenta). x240 actual speed.

**Movie S5.**

Time-lapse during the first cleavage of a starfish embryo expressing POLArIS^act^ with two-axis FPM (green/magenta), related to Fig. 4A. In the first half, cross marks indicate the centers of the green/magenta cross patterns. These illustrations are omitted from the second half. x505 actual speed.

**Movie S6.**

Overnight time-lapse two-axis FPM (green/magenta) after the fourth cleavage of a starfish embryo expressing POLArIS^act^, related to Fig. 4B. Cross marks indicate the centers of the green/magenta cross patterns. x612 actual speed.

**Movie S7.**

Time-lapse green/magenta FPM movie of the first cleavage of a starfish embryo expressing POLArIS^act^ and EMTB-mCherry, related to Fig. 5C. For EMTB-mCherry, only vertical polarization images were shown. In the first half, cross marks indicate the centers of FLARE, and asterisks and dashed lines indicate centrosomes and contours of microtubule asters, respectively. These illustrations are omitted from the second half. x900 actual speed.

## References

1. V. M. Ohmachi, et al., Fluorescence microscopy for simultaneous observation of 3D orientation and movement and its application to quantum rod-tagged myosin V. Proc. Natl. Acad. Sci. U. S. A. 109, 5294–8 (2012).

2. J. N. Forkey, M. E. Quinlan, M. A. Shaw, J. E. T. Corrie, Y. E. Goldman, Three-dimensional structural dynamics of myosin V by single-molecule fluorescence polarization. Nature 422, 399–404 (2003).

3. H. Sosa, E. J. G. Peterman, W. E. Moerner, L. S. B. Goldstein, ADP-induced rocking of the kinesin motor domain revealed by single-molecule fluorescence polarization microscopy. Nat. Struct. Biol. 8, 540–4 (2001).

4. T. Nishizaka, et al., Chemomechanical coupling in F1-ATPase revealed by simultaneous observation of nucleotide kinetics and rotation. Nat. Struct. Mol. Biol. 11, 142–8 (2004).

5. V. Swaminathan, et al., Actin retrograde flow actively aligns and orients ligand-engaged integrins in focal adhesions. Proc. Natl. Acad. Sci. U. S. A. 114, 10648–53 (2017).

6. P. Nordenfelt, et al., Direction of actin flow dictates integrin LFA-1 orientation during leukocyte migration. Nat. Commun. 8, 2047 (2017).

7. S. B. Mehta, et al., Dissection of molecular assembly dynamics by tracking orientation and position of single molecules in live cells. Proc. Natl. Acad. Sci. 113, E6352–E6361 (2016).

8. M. Kampmann, C. E. Atkinson, A. L. Mattheyses, S. M. Simon, Mapping the orientation of nuclear pore proteins in living cells with polarized fluorescence microscopy. Nat. Struct. Mol. Biol. 18, 643–9 (2011).

9. N. Nakai, et al., Genetically encoded orientation probes for F-actin for fluorescence polarization microscopy. Microscopy 68, 359–68 (2019).

10. A. M. Vrabioiu, T. J. Mitchison, Structural insights into yeast septin organization from polarized fluorescence microscopy. Nature 443, 466–9 (2006).

11. A. M. Vrabioiu, T. J. Mitchison, Symmetry of septin hourglass and ring structures. J. Mol. Biol. 372, 37–49 (2007).

12. B. S. DeMay, et al., Septin filaments exhibit a dynamic, paired organization that is conserved from yeast to mammals. J. Cell Biol. 193, 1065–81 (2011).

13. K. Zhanghao, et al., Super-resolution dipole orientation mapping via polarization demodulation. Light Sci. Appl. 5, e16166 (2016).

14. C. A. V. Cruz, et al., Quantitative nanoscale imaging of orientational order in biological filaments by polarized superresolution microscopy. Proc. Natl. Acad. Sci. U. S. A. 113, E820–E828 (2016).

15. A. S. Backer, M. Y. Lee, W. E. Moerner, Enhanced DNA imaging using super-resolution microscopy and simultaneous single-molecule orientation measurements. Optica 3, 3–6 (2016).

16. K. Zhanghao, et al., Super-resolution imaging of fluorescent dipoles via polarized structured illumination microscopy. Nat. Commun. 10, 4694 (2019).

17. R. Bedford, et al., Alternative reagents to antibodies in imaging applications. Biophys. Rev. 9, 299–308 (2017).

18. C. Tiede, et al., Adhiron: A stable and versatile peptide display scaffold for molecular recognition applications. Protein Eng. Des. Sel. 27, 145–55 (2014).

19. D. J. Hughes, et al., Generation of specific inhibitors of SUMO-1- and SUMO-2/3- mediated protein-protein interactions using Affimer (Adhiron) technology. Sci. Signal. 10, eaaj2005 (2017).

20. A. Lopata, et al., Affimer proteins for F-actin: Novel affinity reagents that label F-actin in live and fixed cells. Sci. Rep. 8, 6572 (2018).

21. A. Lopata, et al., Artificial actin-binding proteins with novel multifunctional properties. Biophisical Soc. 60th Annu. Meet., L3598-Pos, Los Angeles, California, 02 Mar (2016).

22. J. Riedl, et al., Lifeact: a versatile marker to visualize F-actin. Nat. Methods 5, 605– 7 (2008).

23. S. J. Winder, et al., Utrophin actin binding domain: analysis of actin binding and cellular targeting. J. Cell Sci. 108, 63–71 (1995).

24. R. Arai, H. Ueda, A. Kitayama, N. Kamiya, T. Nagamune, Design of the linkers which effectively separate domains of a bifunctional fusion protein. Protein Eng. 14, 529–32 (2001).

25. E. F. Pettersen, et al., UCSF Chimera - A visualization system for exploratory research and analysis. J. Comput. Chem. 25, 1605–12 (2004).

26. D. P. Barondeau, C. J. Kassmann, J. A. Tainer, E. D. Getzoff, Understanding GFP posttranslational chemistry: Structures of designed variants that achieve backbone fragmentation, hydrolysis, and decarboxylation. J. Am. Chem. Soc. 128, 4685–93 (2006).

27. K. Brejc, et al., Structural basis for dual excitation and photoisomerization of the Aequorea victoria green fluorescent protein. Proc. Natl. Acad. Sci. U. S. A. 94, 2306–11 (1997).

28. M. Mori, et al., An Arp2/3 nucleated F-actin shell fragments nuclear membranes at nuclear envelope breakdown in starfish oocytes. Curr. Biol. 24, 1421–8 (2014).

29. Y. Hamaguchi, T. Numata, S. K. Satoh, Quantitative analysis of cortical actin filaments during polar body formation in starfish oocytes. Cell Struct. Funct. 32, 29–40 (2007).

30. L. Santella, N. Limatola, F. Vasilev, J. T. Chun, Maturation and fertilization of echinoderm eggs: Role of actin cytoskeleton dynamics. Biochem. Biophys. Res. Commun. 506, 361–71 (2018).

31. M. Mori, et al., Intracellular transport by an anchored homogeneously contracting F-actin meshwork. Curr. Biol. 21, 606–11 (2011).

32. N. Wesolowska, et al., Actin assembly ruptures the nuclear envelope by prying the lamina away from nuclear pores and nuclear membranes in starfish oocytes. eLife 9, e49774 (2020).

33. P. Lénárt, et al., A contractile nuclear actin network drives chromosome congression in oocytes. Nature 436, 812–8 (2005).

34. R. L. Lamason, M. D. Welch, Actin-based motility and cell-to-cell spread of bacterial pathogens. Curr. Opin. Microbiol. 35, 48–57 (2017).

35. K. N. Richter, et al., Glyoxal as an alternative fixative to formaldehyde in immunostaining and super-resolution microscopy. EMBO J. 37, 139–59 (2018).

36. A. L. Miller, W. M. Bement, Regulation of cytokinesis by Rho GTPase flux. Nat. Cell Biol. 11, 71–7 (2009).

37. K. J. Amann, T. D. Pollard, The Arp2/3 complex nucleates actin filament branches from the sides of pre-existing filaments. Nat. Cell Biol. 3, 306–10 (2001).

38. D. Pruyne, et al., Role of formins in actin assembly: Nucleation and barbed-end association. Science 297, 612–5 (2002).

39. B. J. Nolen, et al., Characterization of two classes of small molecule inhibitors of Arp2/3 complex. Nature 460, 1031–4 (2009).

40. S. A. Rizvi, et al., Identification and characterization of a small molecule Inhibitor of formin-mediated actin assembly. Chem. Biol. 16, 1158–68 (2009).

41. S. Terada, T. Nakata, A. C. Peterson, N. Hirokawa, Visualization of slow axonal transport in vivo. Science 273, 784–8 (1996).

42. S. Terada, M. Kinjo, N. Hirokawa, Oligomeric tubulin in large transporting complex is transported via kinesin in squid giant axons. Cell 103, 141–55 (2000).

43. H. F. Kyle, et al., Exploration of the HIF-1α/p300 interface using peptide and Adhiron phage display technologies. Mol. Biosyst. 11, 2738–49 (2015).

44. C. Tiede, et al., Affimer proteins are versatile and renewable affinity reagents. eLife 6, e24903 (2017).

45. K. J. Kearney, et al., Affimer proteins as a tool to modulate fibrinolysis, stabilize the blood clot, and reduce bleeding complications. Blood 133, 1233–44 (2019).

46. E. L. Hesketh, et al., Affimer reagents as tools in diagnosing plant virus diseases. Sci. Rep. 9, 7524 (2019).

47. F. Klont, M. Hadderingh, P. Horvatovich, N. H. T. Ten Hacken, R. Bischoff, Affimers as an alternative to antibodies in an affinity LC-MS assay for quantification of the soluble receptor of advanced glycation end-products (sRAGE) in human serum. J. Proteome Res. 17, 2892–9 (2018).

48. J. I. Robinson, et al., Affimer proteins inhibit immune complex binding to FcγRIIIa with high specificity through competitive and allosteric modes of action. Proc. Natl. Acad. Sci. U. S. A. 115, E72–E81 (2018).

49. M. A. Michel, K. N. Swatek, M. K. Hospenthal, D. Komander, Ubiquitin linkage-specific affimers reveal insights into K6-linked ubiquitin signaling. Mol. Cell 68, 233–246.e5 (2017).

50. S. Inoué, Polarization optical studies of the mitotic spindle. Chromosoma 5, 487– 500 (1953).

51. S. Inoué, Cell division and the mitotic spindle. J. Cell Biol. 91, 131–47 (1981).

52. J. Azoury, et al., Spindle positioning in mouse oocytes relies on a dynamic meshwork of actin filaments. Curr. Biol. 18, 1514–9 (2008).

53. M. Schuh, J. Ellenberg, A new model for asymmetric spindle positioning in mouse oocytes. Curr. Biol. 18, 1986–92 (2008).

54. B. Mogessie, M. Schuh, Actin protects mammalian eggs against chromosome segregation errors. Science 357, eaal1647 (2017).

55. S. Woolner, L. L. O’Brien, C. Wiese, W. M. Bement, Myosin-10 and actin filaments are essential for mitotic spindle function. J. Cell Biol. 182, 77–88 (2008).

56. A. M. Kita, et al., Spindle–F-actin interactions in mitotic spindles in an intact vertebrate epithelium. Mol. Biol. Cell 30, 1645–54 (2019).

57. C. M. Field, J. F. Pelletier, T. J. Mitchison, Disassembly of actin and keratin networks by Aurora B kinase at the midplane of cleaving Xenopus laevis eggs. Curr. Biol. 29, 1999–2008.e4 (2019).

58. R. Rappaport, Experiments concerning the cleavage stimulus in sand dollar eggs. J. Exp. Zool. 148, 81–9 (1961).

59. C. M. Field, A. C. Groen, P. A. Nguyen, T. J. Mitchison, Spindle-to-cortex communication in cleaving, polyspermic Xenopus eggs. Mol. Biol. Cell 26, 3628– 40 (2015).

60. G. von Dassow, Concurrent cues for cytokinetic furrow induction in animal cells. Trends Cell Biol. 19, 165–73 (2009).

61. S. Woolner, W. M. Bement, Unconventional myosins acting unconventionally. Trends Cell Biol. 19, 245–52 (2009).

62. F. Farina, et al., The centrosome is an actin-organizing centre. Nat. Cell Biol. 18, 65–75 (2016).

63. F. Farina, et al., Local actin nucleation tunes centrosomal microtubule nucleation during passage through mitosis. EMBO J. 38, e99843 (2019).

64. D. Inoue, et al., Actin filaments regulate microtubule growth at the centrosome. EMBO J. 38, e99630 (2019).

65. F. Spira, et al., Cytokinesis in vertebrate cells initiates by contraction of an equatorial actomyosin network composed of randomly oriented filaments. eLife 6, e30867 (2017).

66. X. Huang, A. Aulabaugh, Application of fluorescence polarization in HTS assays. Methods Mol. Biol. 1439, 115–30 (2016).

67. A. L. Mattheyses, A. D. Hoppe, D. Axelrod, Polarized fluorescence resonance energy transfer microscopy. Biophys. J. 87, 2787–97 (2004).

68. N. Ojha, K. H. Rainey, G. H. Patterson, Imaging of fluorescence anisotropy during photoswitching provides a simple readout for protein self-association. Nat. Commun. 11, 21 (2020).

69. J. Yamamoto, et al., Rotational diffusion measurements using polarization-dependent fluorescence correlation spectroscopy based on superconducting nanowire single-photon detector. Opt. Express 23, 32633–42 (2015).

70. M. Oura, et al., Polarization-dependent fluorescence correlation spectroscopy for studying structural properties of proteins in living cell. Sci. Rep. 6, 31091 (2016).

71. Y. Zhao, et al., An expanded palette of genetically encoded Ca^2+^ indicators. Science 333, 1888–91 (2011).

72. D. G. Gibson, et al., Enzymatic assembly of DNA molecules up to several hundred kilobases. Nat. Methods 6, 343–5 (2009).

73. W. Kabsch, XDS. Acta Crystallogr. Sect. D 66, 125–32 (2010).

74. A. J. McCoy, et al., Phaser crystallographic software. J. Appl. Crystallogr. 40, 658– 74 (2007).

75. P. D. Adams, et al., PHENIX: A comprehensive Python-based system for macromolecular structure solution. Acta Crystallogr. Sect. D Biol. Crystallogr. 66, 213–21 (2010).

76. P. Emsley, B. Lohkamp, W. G. Scott, K. Cowtan, Features and development of Coot. Acta Crystallogr. Sect. D Biol. Crystallogr. 66, 486–501 (2010).

77. M. D. Winn, et al., Overview of the CCP4 suite and current developments. Acta Crystallogr. Sect. D Biol. Crystallogr. 67, 235–42 (2011).

78. S. Inoué, O. Shimomura, M. Goda, M. Shribak, P. T. Tran, Fluorescence polarization of green fluorescence protein. Proc. Natl. Acad. Sci. U. S. A. 99, 4272– 7 (2002).

79. F. I. Rosell, S. G. Boxer, Polarized absorption spectra of green fluorescent protein single crystals: Transition dipole moment directions. Biochemistry 42, 177–83 (2003).

